# Theory of spontaneous persistent activity and inactivity *in vivo* reveals differential cortico-entorhinal functional connectivity

**DOI:** 10.1101/2022.04.15.488496

**Authors:** Krishna Choudhary, Sven Berberich, Thomas T. G. Hahn, Mayank R. Mehta

## Abstract

Persistent activity is thought to mediate working memory. While such stimulus evoked persistence is well studied, mechanisms of internally generated or spontaneous persistence *in vivo* are unknown. Further, current theories based on attractor dynamics focus on elevated activity as a memory substrate, while little attention has focused on the role of inactivity attractors. Here, we present a mean field model of functional interaction between large cortical networks that predicts both spontaneous persistent activity (SPA) and inactivity (SPI); the latter has never been seen before in experiments or models. We confirm these predictions using simultaneously recorded neocortical local field potential (LFP) and the membrane potential (*V_m_*) of identified excitatory neurons from several brain areas *in vivo* during slow oscillations, especially from layer 3 of the medial (MECIII) and lateral entorhinal cortex (LECIII), which show *SPA* and *SPI*. By matching model and experimental statistics, we predict the relative strength of internal and external excitation in the LECIII and MECIII networks. Our predictions match anatomical data. Further, the model predicts, and the experiments confirm, that *SPA* and *SPI* are quantized by cortical UDS and follow the statistics of a history dependent Bernoulli process. These convergent, theory-experiment results thus reveal the differential nature of cortico-entorhinal functional connectivity, resulting in a unique pattern of persistent activity and persistent inactivity, a novel and energetically efficient memory substrate.

## Introduction

Cognition requires the interaction between several neural networks, each network containing millions of neurons, each neuron in turn characterized by many microscopic parameters. To study the complex emergent properties of systems with large degrees of freedom, the statistical physics approach is to develop a quantitative model, based on only the salient order parameters, and subsequently test its predictions in simplified experimental preparations that capture the essence. In line with this tradition, we develop an analytically tractable model of spontaneous activity in interacting neural networks, and quantitatively verify several predictions of the theory *in vivo* during default, internally generated activity in the absence of external sensory stimuli.

During quiescence, deep sleep, under anesthesia, and *in vitro,* local neural networks from many brain areas, including cortex, show synchronous, rhythmic activity termed delta oscillations, non-REM sleep oscillations, slow wave sleep (SWS) etc.^1–4^. The LFP shows rapid transitions between periods of elevated activity (the Up state) and silence (the Down state). The membrane potential *(V_m_)* of individual neurons exhibits synchronous transitions between depolarized (Up) and hyperpolarized (Down) states. These Up-Down states (UDS) are ubiquitously found across species and experimental preparations, and are considered the default activity of many networks^5–8^. Several studies have suggested that the interactions between cortical regions during UDS are crucial for memory consolidation^9–13^. Impairment of UDS causes learning and memory deficits, while UDS enhancement leads to improvement^14,15^.

Although most cortical areas show synchronous UDS oscillations^11^, recent studies have shown that *in vivo* only MECIII, but not LECIII, pyramidal neurons show spontaneous persistent activity *(SPA)* during UDS: events where the neuron’s *V_m_* persists in the depolarized Up state while the afferent neocortical areas transition to the Down state^16^. This definition notably differs from other studies that define singular Up states within an isolated network as themselves forms of persistent activity^17^. Instead, it is reminiscent of activity sustained after the extinguishing of a stimulus, hypothesized to form the neural representation for working memory during awake behavior. Existing network models show that such sustained activity during awake, working memory tasks can be generated through reverberant excitation and feedback inhibition, but it is unclear whether these models can explain spontaneously evoked persistent activity^18,19^. Depolarizing current injections do not elicit *SPA* within MECIII neurons during UDS, implicating network rather than intracellular mechanisms^16^. Existing network models of UDS employ an attractor framework with two fixed points, one active (the Up state) and one inactive (the Down state), with adaptation driving the oscillation^20–24^. Such models, however, have not been used to understand large interacting networks, and thus cannot account for major experimental findings, like the quantization of SPA during UDS^25^. Furthermore, existing theories focus exclusively on the active state, discarding the inactive state as simply a recovery phase for network adaptation. The energy function of discrete Hopfield networks^26,27^, however, is symmetric under activity inversion (+1 → −1), so the physics suggests these inactive states are themselves energy minima in the landscape and could thus also be utilized as a memory substrate.

We found that a simple, mean-field model involving two interacting networks of excitation-inhibition can capture the dynamics of *SPA* during UDS. Our theory also exploited the symmetric inactive attractor to predict a new phenomenon: spontaneous persistent inactivity *(SPI).* To test the model quantitatively, we used the *in vivo* cortico-entorhinal circuit as our model system. Anatomically, the neocortex serves as an afferent source of input to other cortical regions like the entorhinal cortex^28,29^. To measure neocortical ensemble activity during UDS, we recorded the extracellular LFP from the parietal cortex. As the parietal cortex receives strong inputs from neocortical regions^30–33^ and UDS is synchronous across all neocortical areas^2,11,34,35^ this LFP acted as the afferent reference for neocortical UDS. Simultaneously, we did wholecell *V_m_* measurements from anatomically identified pyramidal neurons in various efferent areas, including parietal (PAR) and entorhinal cortices (EC); as the spontaneous activity of single neurons is tightly linked to the cortical networks in which they are embedded, this allowed us to probe the activity of localized networks within each target region^36^. Within the EC, the medial (MEC) and lateral (LEC) subdivisions are anatomically and functionally distinct: the MEC contains spatially selective “grid cells”^37^, while the LEC is thought to encode objects or experienced time^38–43^. We focused in particular on the EC layer 3 regions, since MECIII neurons are a major source of input to the hippocampus, show *SPA in vivo,* and are crucial in the generation and maintenance of UDS in the MEC^16,44^.

We detected *in vivo SPA* in MECIII and *SPI* in both MECIII and LECIII, but not in PAR. Further analysis of these events showed clear agreement with theoretical predictions, as both *SPA* and *SPI* were quantized by neocortical UDS and reflected the statistics of a history-dependent Bernoulli process. Our model attributed the differences in *SPA* and *SPI* across cells to differences in excitatory connectivity between the large neocortical network and the specific efferent subnetwork. The number of experimental observations explained by our model are greater than the number of parameters we varied, demonstrating its predictive power. To our knowledge, our study is the first to predict theoretically and detect experimentally the novel phenomena of persistent inactivity, and show that both *SPA* and *SPI* not only cooccur but are the result of common network interaction principles.

### The mean field model of cortical interaction predicts both spontaneous persistent activity *(SPA)* and inactivity *(SPI)*

A minimal mean field network supporting UDS has three biologically well-established ingredients: excitation, inhibition, and the adaptation of excitation (but not inhibition)^20–22,24^. We constructed a mean field model of two cortical regions, each with its own recurrently connected inhibitory and excitatory populations^45,46^ (Fig 1). In isolation, each network exhibits transitions between Up and Down states (Sup. Fig 1-2) that are the stable fixed points of the dynamical system of equations, much like local minima in an energy landscape^47,48^. Their stability is inversely related to their distance from the separatrix, a line which defines the boundary between the basins of attraction. The slowly-varying, activity-dependent adaptation translates the excitatory nullcline, thus influencing the stability of each state. Growing adaptation governs the transition from the Up to Down state, while external drive and a falling adaptation governs the transition from Down to Up. Underlying gaussian noise gives the network a “temperature,” preventing it from stagnating in a particular state for arbitrarily long time periods. For simplicity, quantitative falsifiability, and based on available observations, we assumed that all internal parameters except the recurrent excitation strength *W*_INT_ are identical across the afferent and efferent networks.

**Figure 1:**
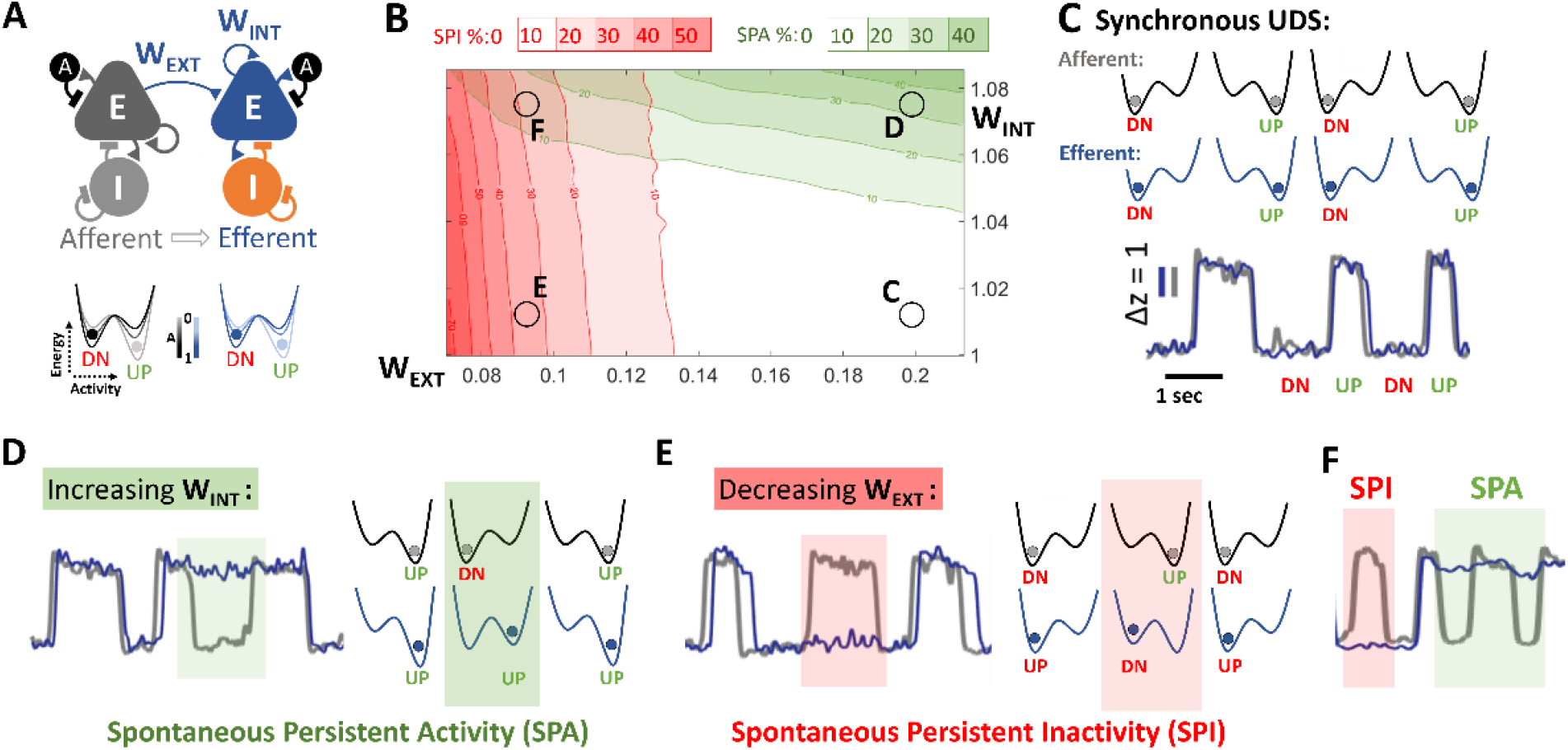
A simple mean-field model predicts *spontaneous persistent activity* and *inactivity* in the efferent network. **A**) The model consists of two networks, each characterized by the average activity of excitatory *(E)* and inhibitory *(I)* populations, with only the *E* showing activity dependent adaptation (A). Each network is described by a potential landscape with two local minima, the Up and Down states. The changing adaptation level influences the stability of each minima, creating the Up-Down state (UDS) oscillation. The afferent network provides excitatory input (*W_EXT_)* to the efferent network. All internal parameters are identical between the two networks, except *W_INT_*, the recurrent excitation. **B)** By modulating *W_INT_* and *W_EXT_*, the model introduces transient desynchronizations between the two networks. There are four distinct regimes (red and blue shaded areas, C, D, E, F): **C)** The model can reproduce synchronized UDS in the two networks (afferent in gray, efferent in blue). The same scale bars for amplitude (in z-score) and time (1 sec) are used for (D-F). **D)** Increasing efferent *W_INT_* increases the stability of the efferent Up state attractor, and leads to *spontaneous persistent activity (SPA,* green box), when the efferent network gets “stuck” or persists in the Up state even when the afferent makes a transition to the Down state. Green contours in (B) show SPA levels in the parameter space. **E)** Conversely, decreasing *W_EXT_* decreases the size of destabilizing afferent current and leads to *spontaneous persistent inactivity (SPI,* red box), when the efferent network persists in the Down state while the afferent makes a transition to Up state. Red contours in (B) show SPI levels. **F**) The same network can exhibit *SPA* and *SPI* at different times. The positions of each example in the 2D parameter plane is shown in B.

These two networks are connected unidirectionally, with the afferent network sending an excitatory projection *W_EXT_* to the excitatory population in the efferent network (Fig 1A, Sup. Fig 3). The UDS oscillation of the afferent network rhythmically destabilizes the endogenous UDS oscillation of the efferent network. While larger values of *W_EXT_* lead to phase locking (Fig 1C), smaller values give rise to transient desynchronizations between the two networks^49^. Under weak drive *W_EXT_* from the afferent network, an increase in the efferent recurrent excitatory connectivity (*W_INT_*) drives the efferent Up state fixed point away from the separatrix, increasing the Up state stability. Simulations demonstrate a novel, network mechanism to generate *SPA*: instances when the efferent network remains in the Up state, skipping one or more afferent Down states (Fig 1D). These could explain the experimentally reported *SPA*. Further, a decrease in the strength of *W_EXT_* decreases the destabilizing effect of the afferent transitions on the efferent network; the efferent then remains in a Down state, skipping one or more afferent Up states. We call this novel phenomena *spontaneous persistent inactivity (SPI:* Fig 1E). The model further predicts that *SPA* and *SPI* are relatively independent, as increasing *W_INT_* while simultaneously decreasing *W_EXT_* gives rise to coupled UDS sequences exhibiting both *SPA* and *SPI* (Fig 1F).

### Detection of *SPA* and *SPI* in MECIII and LECIII

To monitor network interactions during spontaneous activity *in vivo,* mice were lightly anesthetized with urethane to induce robust and steady UDS that were synchronous across the entire neocortex. A hidden Markov model was used to classify the data into a binary UDS sequence^50^. Consistent with previous studies, the neocortical LFP and the *V_m_* of neurons in PAR (N=24) and the efferent regions MECIII (N=50) and LECIII (N=16) showed clear bimodal UDS (Fig 2, Sup. Fig. 4). For subsequent analysis, the amount of *SPA (SPI)* was defined as the proportion of efferent Up (Down) states which outlasted an entire afferent Down (Up) state during an entire experiment. As a first test of the model, we computed the relationship between the neocortical LFP and the *V_m_* from PAR pyramidal neurons, which were recorded close by (0.5 mm apart). Here, *W_EXT_* is large, and the model predicts complete phase locking (Fig 1C), with virtually nonexistent *SPA* and *SPI*. This was indeed the case (Fig 2B). Additional *Vm* measurements from neurons in frontal and prefrontal cortex also showed complete phase locking, consistent with *W_EXT_* from the neocortex being large (Sup. Fig. 4)^11,30–35^.

**Figure 2:**
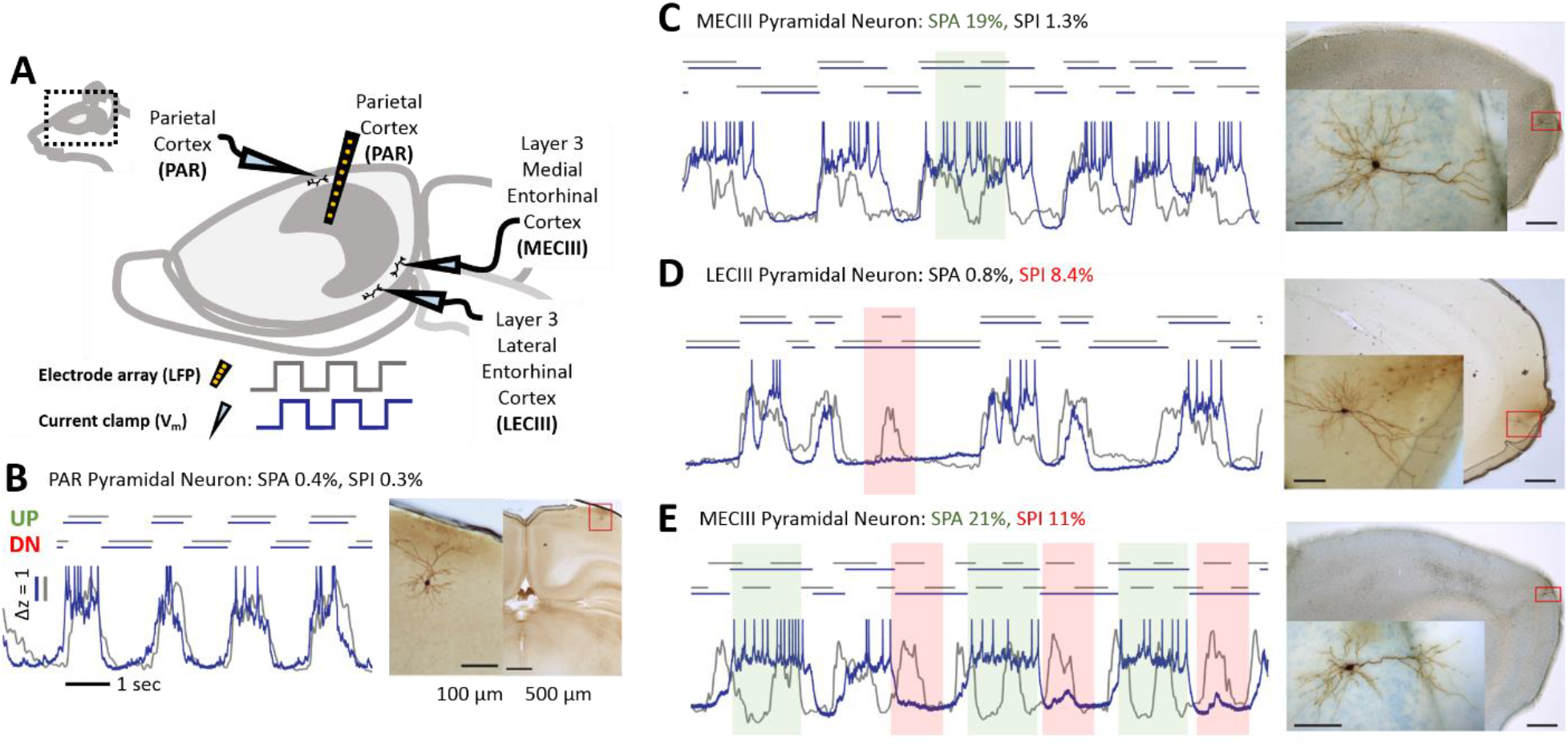
An *in vivo* study of simultaneous UDS in neocortical and entorhinal networks confirms prediction of SPA and SPI. **A**) Experimental design: mice were anesthetized to induce UDS, and local field potential (LFP) from the parietal cortex was measured using a silicon probe (black). Simultaneously, membrane potential (*V_m_*) was measured using whole-cell patch clamp from an anatomically identified neuron. The parietal LFP is treated as the afferent reference representing neocortical activity (gray in B-E), and the *V_m_* traces are the efferent activity (blue in B-E). **B)** PAR neuron’s *V_m_* is phase-locked to the neocortical LFP, matching theory (Fig. 1C). Action potentials have been truncated for clarity. The same scale bars for time (1 second) and amplitude (z-scores) are used throughout. The identified UDS sequence is shown above the traces, with histological reconstructions (right) of brain region and the patched cell (insets). **C)** Clear *SPA* (green box) in the *V_m_* of an MECIII pyramidal neuron, matching (Fig. 1D). **D)** Clear *SPI* (red box) in the *V_m_* of an LECIII pyramidal neuron, matching (Fig. 1E). **E)** Both *SPA* and *SPI* at different times exhibited by the same MECIII pyramidal neuron, similar to (Fig. 1F).

Consistent with previous studies, MECIII neurons showed clear instances of *SPA* (Fig 2C), while LECIII neurons did not^16^. In contrast, both LECIII and MECIII neurons showed clear instances of the newly predicted *SPI* (Fig 2D). Our model also predicted relative independence of *SPA* and *SPI*; consistently, some MECIII neurons showed both *SPA* and *SPI*, only a few seconds apart (Fig 2F), and levels of *SPA* and *SPI* within the population of LECIII and MECIII neurons were not significantly correlated (Sup. Fig. 5). Finally, *SPA* and *SPI* levels were not correlated with the duty cycle and the frequency of neocortical UDS, indicating that they were not artifacts of differences in brain states across experiments (Sup. Fig. 6).

### Fitting experiment to model reveals differential connectivity within MECIII and LECIII

The properties of *SPA* and *SPI* not only varied across brain regions, but even between different neurons from the same region. We hypothesized that all of these differences could arise from just two network parameters: the strength of recurrent excitation in the efferent network (*W_INT_*) and the strength of external excitatory input to the efferent network (*W_EXT_*). To test this idea, we used a two-step approach. First, we simulated all possible networks in this 2D parameter space by varying only *W_EXT_* and *W_INT_*, while leaving all other variables unchanged. Modulating just two free parameters yielded networks with a wide range of both *SPA* and *SPI*. Thus, we could estimate the two crucial network variables, *W_INT_* and *W_EXT_*, by simply computing the amount of *SPA* and *SPI* observed experimentally (Fig 3A). Crucially, we did not match any other properties of *SPA* and *SPI* between the model and data. The robustness of this procedure was confirmed by using an alternate fitting procedure, which yielded very similar fits between the simulations and *in vivo* data for each neuron (Sup. Fig 7). Overall, a decrease in *W_EXT_* corresponded with higher levels of *SPI*, and an increase in *W_INT_* corresponded with higher *SPA*, as predicted by dynamical systems analysis (Fig 3A, B). While *SPA* and *SPI* prevalence across neurons was uncorrelated, the fitted values of *W_INT_* and *W_EXT_* were significantly negatively correlated, especially for LECIII, indicating differential properties of the networks (Sup. Fig 5).

**Figure 3:**
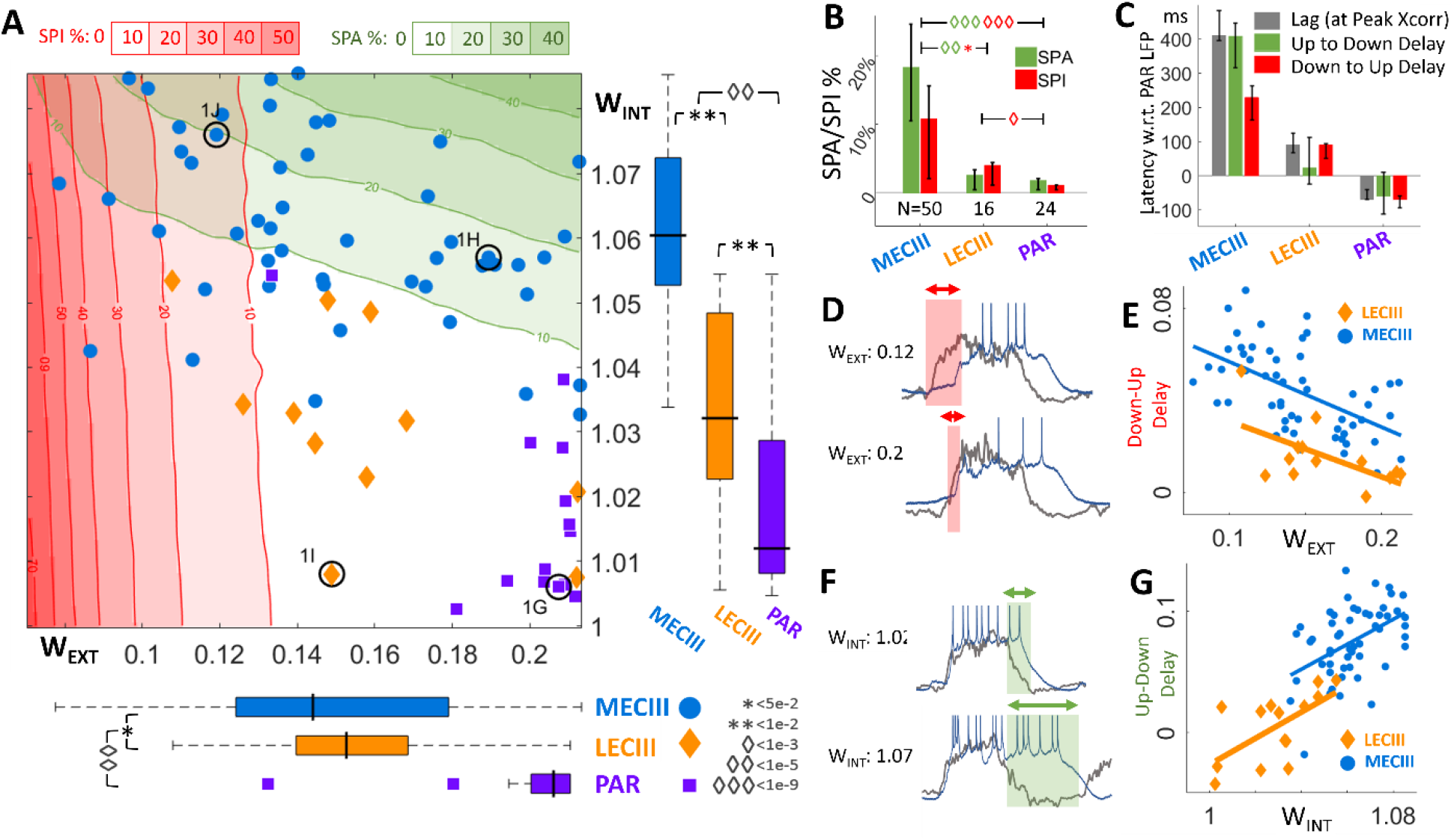
Model predicts larger transition delays in MECIII than LECIII based on differences in the strength of recurrent (*W_INT_*) and external (*W_EXT_*) inputs: **A**) The time-averaged prevalence of *SPA*% (green) and *SPI*% (red) observed in a single cell were reproduced in the model by varying just two variables, *W_EXT_* (x-axis) from the neocortex and *W_INT_* (y-axis) within the efferent subnetwork. The contour lines and shaded areas reveal that *W_INT_* has a greater impact on *SPA* (nearly horizontal green lines), whereas *W_EXT_* has a greater impact on *SPI* (nearly vertical red lines). Each cell is represented by a point (blue circle MECIII, yellow diamond LECIII, violet square parietal (PAR)). Locations of the examples cells used in Fig. 2 are highlighted by the black circles. Bottom: The fitting reveals that *W_EXT_* is the largest for PAR (violet, 0.21±0.01), smaller for LECIII (yellow, 0.15±0.02), and smallest for MECIII (blue, 0.14±0.03). *W_EXT_* to MECIII was significantly smaller than to LECIII (p<0.05), and both were significantly smaller than to PAR (p<10^-5^). Box limits and black bar represents middle 50% of data and median, and dotted lines extend to the range, except outliers. Right: The fitting predicts that *W_INT_* is the largest for MECIII (1.06±0.01), smaller for LECIII (1.03±0.015), and even smaller for PAR (1.01±0.02). MECIII *W_INT_* was significantly larger than LECIII *W_INT_* (p<10^-2^), and both were larger than in PAR (p<10^-2^). **B**) Prevalence of *SPA* and *SPI* showed significant variation as a function of brain region, with MECIII showing the highest *SPA* (18.5±7.5%) as well as *SPI* (10.2±5.3%). LECIII showed diminished *SPA* (3.3±2.4%) and *SPI* (5.9±3.1%), while PAR *(SPA:* 2.6±1.8%, *SPI*: 1.9±0.7%) showed even smaller levels. MECIII *SPA* and *SPI* levels were both significantly different from both LECIII levels (*SPA*: p<10^-5^, *SPI*: p<0.05) and PAR levels (*SPA*: p<10^-9^, *SPI*: p<10^-9^), while for LECIII only the *SPI* level was significantly larger from PAR (p<10^-5^). Error bars represent ±1 s.t.d, and number of experiments within each brain region is indicated under each bar. **C**) Strength of *W_EXT_* predicts the latency of efferent responses, with greater *W_EXT_* resulting in shorter response latency. The model predicts that *W_EXT_* is the smallest for MECIII, and consistently the experimentally observed latency between the neocortical LFP and efferent MECIII cells are the largest, evidenced by cross-correlation latency (gray, 152±145ms), Up-Down latency (green, 401±23ms) and Down-Up latency (red, 223±34ms). The delays for LECIII cells was shorter than those for MECIII cells (Xcorr latency 89±22 ms, Up-Down 22±12 ms, and Down-Up 76±13 ms). PAR cells were unique in that they preceded PAR LFP (Xcorr latency −54±21 ms, Up-Down −43±34 ms, and Down-Up −58±13 ms). **D-G**) The model predicts not only the pattern of latency across brain regions, but difference in latency of individual cells, based on the decoded value of *W_EXT_* and *W_INT_* **D**) An increase in the external excitation *W_EXT_* increases the coupling of the efferent network to the input, decreasing the latency between the afferent and efferent networks. Two example experiments are shown; the cell with larger *W_INT_* shows smaller Down-Up transition delay (red boxes). **E**) The average Down-Up transition latency (normalized by UDS duration) in the experiment was significantly anti-correlated with the predicted value of *W_INT_* for MECIII (blue, r=-0.56, p<10^-5^) and LECIII (yellow, r=-0.60, p<10^-2^). **F**) Larger internal excitation *W_INT_* increases the stability of the Up attractor in the efferent network, resulting in longer persistence in the Up state. Examples from two experiments show the cell with larger predicted value of *W_INT_* having greater Up-Down transition lag (green boxes). **G**) The average Up-Down transition latency (normalized by the mean duration of UDS) in the experiment was significantly positively correlated with the predicted value of *W_INT_* for MECIII (blue, r=0.47, p<10^-3^) and LECIII (yellow, r=0.63, p<10^-2^).

The two parameter model, thus constrained by experiments, made major predictions about the nature of large-scale connections between and within these brain regions. Briefly, our model implies that neocortical input into MECIII is weaker than into LECIII, and still weaker than into other neocortical regions, like parietal, frontal, and prefrontal cortices. Further, it predicts that recurrent excitation within MECIII is stronger than within LECIII. These statements are corroborated by established experiments *in vivo* and *in vitro* (see Discussion). Additionally, several further predictions of the model could be tested using the match between experiment and simulation.

### Inferred network connectivity predicts differential latency to UDS transitions in MECIII and LECIII

Since neurons behave like leaky capacitors, the strength of afferent excitatory input should be inversely correlated with the response latency of the efferent neurons^51,52^. Therefore, the model predicts that the neurons with larger values of estimated *W_EXT_* should respond sooner to neocortical Down-Up transitions, i.e. smaller latency between neocortical LFP and the neuron’s *V_m_* (Fig 3C). Indeed, LECIII cells with greater predicted excitatory input *W_EXT_* showed significantly shorter Down-Up transition latency (Fig 3D-E). A similar result was found within the population of MECIII neurons. Further, consistent with model prediction that *W_EXT_* from the neocortex to LECIII is stronger than to MECIII, the population of LECIII neurons showed shorter Down-Up latency than the MECIII population (Fig 3D-E). While *W_EXT_* enhances the coupling between the two networks, larger values of *W_INT_* make the efferent network more independent of the input. The effect of these competing inputs is state dependent, differentially modulating the efferent Down-Up vs. Up-Down transitions. During an afferent Down-Up transition, the efferent network is in the Down state, where recurrent excitation *W_INT_* does not contribute. Thus, the latency of the efferent Down-Up transition should be relatively insensitive to *W_INT_* but depend strongly on *W_EXT_*. This was strongly supported across both LECIII and MECIII populations (Sup. Fig. 8).

The situation is reversed for the Up-Down transition, when the efferent network is in the Up state, where recurrent excitation *W_INT_* contributes strongly and helps sustain the Up state despite the loss of afferent input, which is in the Down state. Networks with higher *W_INT_* have more stable Up states, thereby increasing their “inertia.” Thus, the model predicts that ECIII neurons with greater predicted *W_INT_* should follow the neocortical Up-Down transitions with longer latency. This was confirmed for both MECIIII and LECIII (Fig 3F-G). In contrast to Down-Up transitions, the latency of the efferent Up-Down transition should be relatively insensitive to *W_EXT_* compared to *W_INT_*. This prediction too was supported across individual neurons within MECIII, within LECIII, and across the MECIII vs LECIII ensemble (Sup. Fig. 8).

These latencies were more correlated with the predicted *W_INT_* and *W_EXT_* values than with simply the levels of *SPA* or *SPI* (Sup. Fig. 8), further supporting the model. Additionally, neurons with greater net excitatory input (*W_EXT_* + *W_INT_*) should have higher firing rate; this was confirmed by experiments, showing greater mean firing rates for MECIII than LECIII, even at the level of individual cells (Sup. Fig. 9). Further, LECIII neurons’ *V_m_* was significantly less depolarized than MECIII neurons (Sup. Fig. 9). The predicted model parameters *W_EXT_* and *W_INT_* were more strongly correlated with the UDS latencies than with the mean firing rates, further supporting the model and ruling out nonspecific effects.

### *SPA* and *SPI* are quantized by neocortical UDS

The model predicts that *SPA* and *SPI* are all-or-none events that are initiated and terminated by state transitions in the afferent network. As a result, even though efferent Up(Down) state and *SPA(SPI)* durations form a continuous, unimodal distribution, these durations should be quantized in integral units of the afferent UDS cycles (Fig 4A, Sup. Fig. 10). To visualize this for *SPA*, segments of the simulated efferent activity were extracted around each efferent Down-Up transition, sorted according to the ensuing Up state duration, and assembled into a single matrix, with each row corresponding to a single efferent Down-Up transition (Fig 4B). The underlying afferent activity matrix for the same time points exhibited alternating bands of UDS, with integer multiples of afferent UDS fitting inside each efferent Up state (Fig 4C). The same visualization with *in vivo* data matched strikingly well with model predictions (Fig 4D). We repeated this for efferent Down states and *SPI*, yielding a similar quantitative match between the model and experiment (Fig 4E-F). When consolidating the rescaled state durations over all experiments and their matched simulations, the probability distributions for both were significantly multimodal, with peaks at half integers, indicating that ECIII state transitions were locked to the neocortical transitions, and that the ECIII skipped entire neocortical Up/Down states in integer quantities (Fig 4G, Sup. Fig. 11).

**Figure 4:**
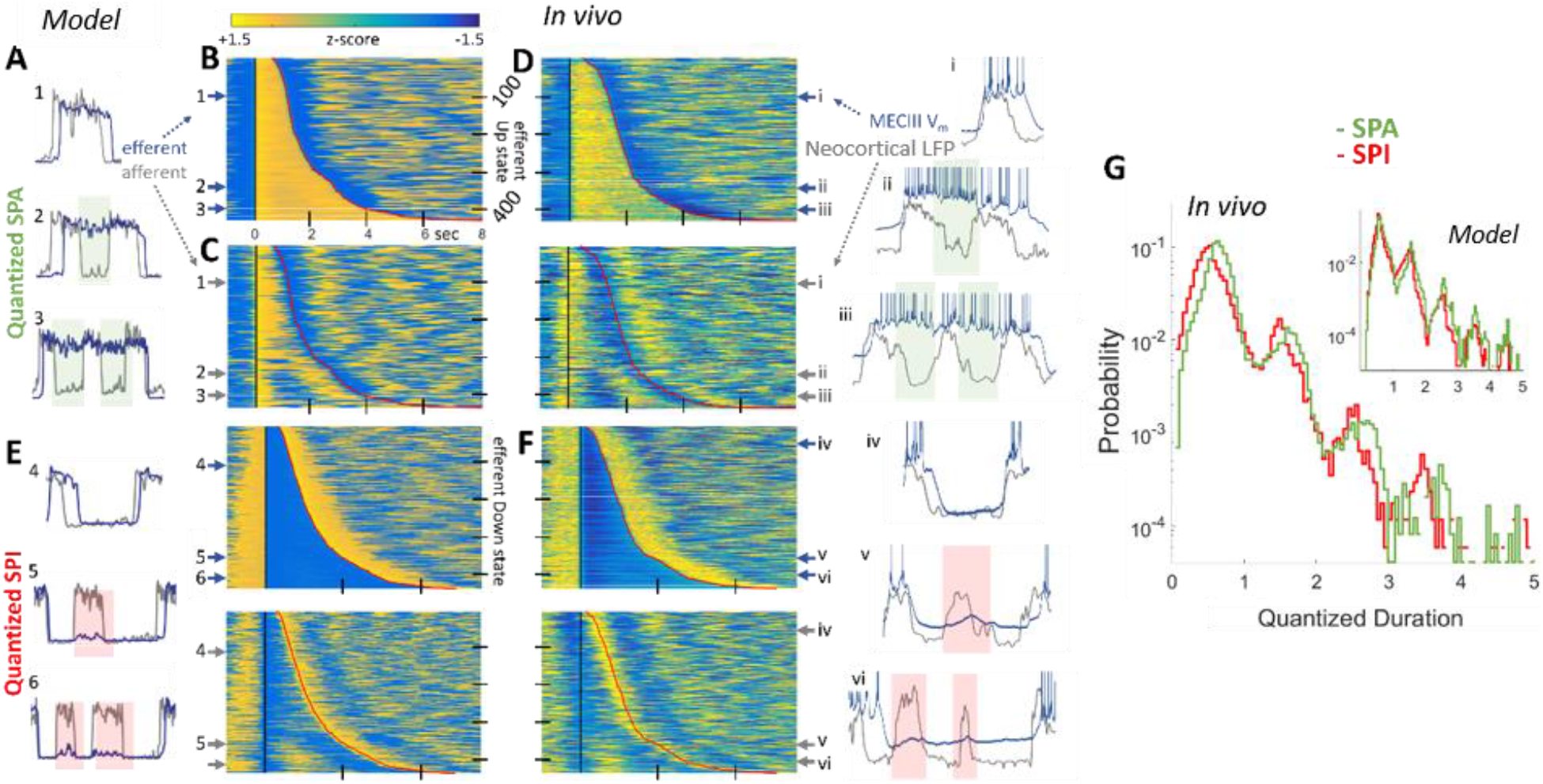
*SPA* and *SPI*are quantized in units of afferent neocortical UDS cycles. **A**) In the model, the efferent *SPA* (blue) span a continuous range of durations, but they are quantized (green boxes, 1,2,3... n) in the units of the number of afferent UDS cycles (gray). **B**) All efferent Up states were aligned to the efferent Down-Up transition and ordered with increasing duration. Each row represents a single efferent Down-Up transition. **C**) A second matrix was constructed using the same time points, but using the afferent network activity. The examples (1-3) in (A) correspond to the row numbers in (B/C). Efferent *SPA* can last 8 seconds, spanning several afferent UDS cycles. Same colorbar axis is used for the amplitude (z-score) for all the panels. **D**) *In vivo* data from an MECIII neuron validates the model by showing efferent MECIII Up states (blue) last an integer multiple (i-iii) of afferent neocortical LFP UDS cycles (gray). **E**) Similar to A-D, but for *SPI*, showing that efferent Down states and *SPI* show a continuous duration but are quantized (red boxes) in the units of afferent neocortical LFP UDS cycles, in the model and **F)** experiment. **G)** With time measured in units of the varying afferent UDS cycles, the model (inset) predicted that the distribution of both Up and Down state durations should be multimodal, with peaks at the half integers (reflecting state transitions after an Up/Down state). *In vivo* data combined from all experiments showed the predicted multimodality and quantization when ECIII Up/Down state durations were measured w.r.t. variable neocortical UDS cycle lengths.

The multimodality of quantized durations was also observed for individual experiments and their corresponding simulations (Fig 5A, Sup. Fig. 11). We leveraged this distribution, unique to each cell, to investigate the precise history-dependence of *SPA* and *SPI*, further testing our model. One can imagine three scenarios. First, the *SPA* and *SPI* are entirely stochastic, in which case their probability distribution would follow a memoryless Bernoulli process, like a sequence of coin flips. Second, *SPA* and *SPI* arise due to some change in the overall state of the animal, such that all the *SPA* and *SPI* co-occur. However, our model predicts a third possibility: it should be rarer to have consecutive sets of *SPA* and *SPI* compared to singular events. This is because the probability of *SPA* and *SPI* is strongly history-dependent. If the network exhibits *SPA* at a given afferent Down state, the efferent network’s recurrent excitation *W_INT_* would be more adapted than usual, reducing the resources needed to sustain *SPA* in the next Down state, thus reducing the probability of consecutive *SPAs*. Similarly, the occurrence of *SPI* at a given afferent Up state would make the efferent network less adapted and hence reduce the probability of consecutive *SPIs*. To test this prediction, we used the first two modes of the quantized probability distribution (in Fig 5A) to calculate *a_1_*, the probability of a solitary *SPA* and *SPI*, and *a_2_*, the probability that another *SPA* and *SPI* occurred given *a*_1_ already happened. Here, *a_2_=a_1_* for the first memoryless hypothesis, *a_2_* > *a_1_* for the second brain-state dependent hypothesis, and *a_1_* > *a_2_* for the third hypothesis, predicted by our model. The experiments strongly corroborated our predictions: the probability of *SPA* and *SPI* diminished after the first such event (Fig 5B). Thus the two network system has a “memory” of *SPA* and *SPI* due to the adaptation of the recurrent excitation *W_INT_* in the efferent network.

**Figure 5:**
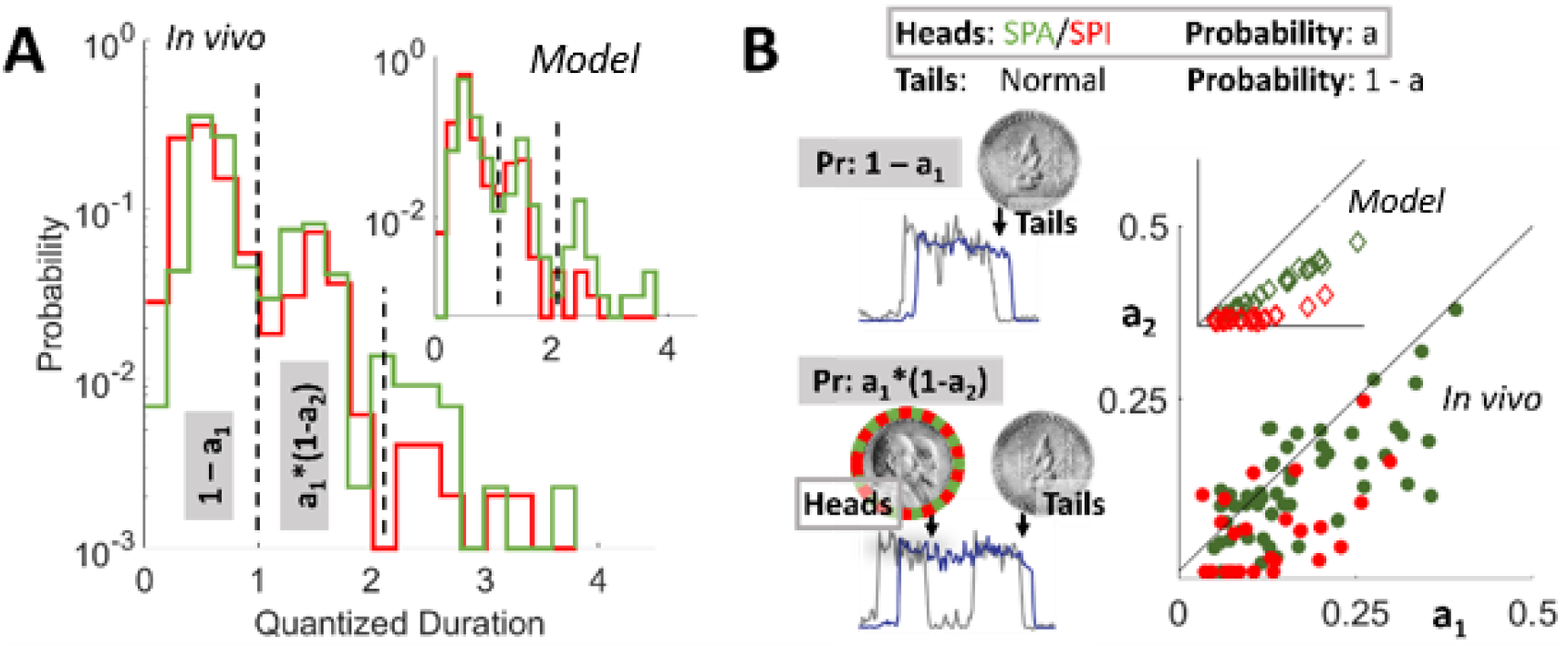
Adaptation introduces history-dependence to SPA and SPI, a Bernoulli process. **A)** Quantization of SPA and SPI is evident within an individual experiment (same as Fig 4) and its corresponding simulation (inset). The probability decreases with SPA and SPI duration, reflecting a discrete Bernoulli process. **B)** To find the precise history-dependence, one can imagine the efferent response to afferent UDS as a sequence of coin flips, as *SPA* and *SPI* are all-or-nothing binary events. State transitions in the afferent network destabilize the efferent network, which either persists in its current state, resulting in *SPA* or *SPI* (heads, probability Pr=a), or makes a corresponding transition (tails, Pr=1-a). The probability that a given efferent state lasts a single afferent UDS cycle is Pr=1-a_1_ (the first mode in G), while the probability that it lasts between 1 and 2 cycles is Pr=a_1_(1-a_2_) (the second mode in G), where the subscript denotes the n^th^ transition. Unlike a memoryless process (i.e. a_1_=a_2_) or brain state dependent process (i.e. a_1_<a_2_), the model (inset) predicted that the probability of *SPA* or *SPI* should decrease when conditioned on *SPA* or *SPI* having occurred previously (i.e. a_1_>a_2_, p<10^-16^). This was confirmed in the *in vivo* data (main, p<10^-7^) and reflects the underlying long-term memory of the adaptation in the efferent network.

## Discussion

Persistent activity has been hypothesized to mediate working memory via reverberating activity^26,53^, and has been studied extensively *in vivo*^54–56^, *in vitro*^57–60^, and *in silica*^18,19^. Its ubiquity and diversity in different cell types, brain regions, brain states, and behavior supports the hypothesis that a common mechanism could apply, and a low dimensional theory could be well suited to explain it. We developed a mean field model to explain the recent discovery of spontaneous persistent activity in MECIII during sleep^16^, as existing models focus only on stimulus evoked persistent activity during awake behavior. Using two networks of excitation-inhibition neurons and adapting excitation, with an afferent network providing excitatory input to an efferent one, our model reproduced phase locked Up-Down state (UDS) oscillations and the reported spontaneous persistent activity.

Further, the model exploited the symmetry of the discrete attractor landscape to make a surprising prediction, namely the existence of persistent inactivity. In contrast to persistent activity, which involves the efferent network sustaining activity while afferent inputs have shut off, persistent inactivity involves the efferent network sustaining *inactivity* while afferent input turns on. This has not been reported before in any experimental or theoretical studies, though there are hints^61,62^. Computational studies have found coexisting Up and Down states in different neurons within the same spiking network^63,64^, but these results are usually achieved when the network UDS is highly irregular and asynchronous. To test our model, we focused on the cortico-entorhinal interaction during UDS oscillations, using simultaneous LFP from the neocortex that served as a common afferent reference, along with the membrane potentials measured from anatomically identified neurons in the parietal, frontal, prefrontal, and entorhinal cortices.

The experiments confirmed the presence of both persistent activity and inactivity; we were then able to leverage these two observables to probe the underlying network architecture. Our framework models different brain regions by varying only two biologically relevant parameters: the strength of internal connections *W_INT_* within the efferent network and the strength of external input *W_EXT_* from the neocortex while leaving all the other parameters unchanged. Dynamical systems analysis^47,48^ showed that SPA increases with *W_INT_*, while SPI decreases with *W_EXT_*; thus, each cell, and the local network in which it is embedded^36^, could be mapped to the *W_INT_-W_EXT_* parameter space. Our results predicted that neocortical input onto the entorhinal region should be weaker than to other regions within the neocortex, like parietal, frontal, and prefrontal cortex. This is consistent with anatomical observations of strong intra-neocortical connections and weaker neocortical-entorhinal connections^29,30,32,65,66^. Within the entorhinal region, the model predicted that neocortical input into LECIII was significantly stronger than to MECIII. This is consistent with classic anatomical studies^67,68^, which show that a higher proportion of LEC afferents originate in cortical areas compared to MEC afferents, and more recent work^40,69^ showing stronger projections from the orbitofrontal cortex, part of the prefrontal cortex, to LEC compared with MEC.

Our analysis found greater amounts of persistent activity in MECIII than LECIII, and the model predicted that this is because the recurrent connections *W_INT_* should be larger within MECIII than within LECIII. This is indirectly supported by recent experiments showing greater recurrent connectivity between principal neurons within MECIII than within other MEC layers, and that MECIII network is crucial in the initiation and maintenance of the Up state during UDS *in vitro* in isolated EC slices^44,70^. Furthermore, excitatory cholinergic receptors are crucial for MECIII persistent activity^71^, and the application of acetylcholine to MEC slice preparations *in vitro* causes prolonged Up states in individual cells due to increased overall excitation and more frequent and rhythmic population-wide events, consistent with our hypothesis that persistent Up states are the result of networks having increased internal excitation *W_INT_*^72^.

The model with above network connectivity not only predicted the prevalence of *SPA* and *SPI* in the efferent neurons but also predicted their relative timing to afferent neocortical activity^51^, at both population-wide and single-cell resolution. Cells with higher predicted *W_EXT_*, and thus stronger coupling, exhibited significantly shorter state transition lags, while larger recurrent excitation *W_INT_*, and thus stronger “inertia,” had longer lags, as expected. The latency patterns were quite different for Down-Up vs. Up-Down transitions: the former was more dependent on *W_EXT_*, and the latter more on *W_INT_*. Our results thus support the hypothesis that the Up state is terminated by internal network mechanisms but is initiated by external input^20,73^. Taken together, these mechanisms resulted in systematic differences in the response latencies of MECIII and LECIII neurons during Up and Down states, which would influence the information processing in downstream hippocampal neurons^11,16^ and hence the memory consolidation process via spike timing-dependent plasticity mechanisms^13^.

As a direct consequence of the underlying physics of the model, we predicted that both *SPA* and *SPI* durations, while showing continuous, long-tailed distributions, should also show quantization in the units of afferent neocortical UDS cycles. This too was verified experimentally, with not just qualitative but a quantitative match between the model and experiment. Our model went further to predict that *SPA* and *SPI* were highly history-dependent, reducing the probability of consecutive *SPA* and *SPI*, and this too was confirmed *in vivo.* This long time-scale memory is an emergent property of the adaptation in the efferent EC network, which has been implicated in the formation and maintenance of periodic spatial firing of grid cells in MEC^74^.

While persistent activity has been studied extensively as the mechanism underlying working memory, it is far more energetically expensive than persistent inactivity. Furthermore, the models involving only persistent activity have a limited storage capacity, especially when dealing with memories that require overlapping representations^75–77^. Persistent inactivity introduces a new mechanism to overcome this difficulty. From an information theoretic perspective, a 0 is just as informative as a 1. Hence, a combination of persistent activity and inactivity would be an energy and information efficient scheme for storing overlapping memories by multiplexing the representation^78,79^. Related, our model predicted that the same neuron can show *SPA* and *SPI*, and this was experimentally confirmed. Recent theories investigated “persistent activity-silent” mechanisms for working memory and hypothesized that the information is stored in facilitated synapses^80–82^. One prediction is that non-specific inputs can reawaken the memory ensemble after the inactive period. Our model predicts, and experiments confirm, something similar: that the efferent network is more susceptible to inputs after SPI due to falling adaptation. These dynamics between adaptation and activity could drive the production of sequences of memories in neural networks with discrete^83^ and continuous phase spaces^84^.

The long duration of UDS under anesthesia allowed unequivocal detection of both *SPA* and *SPI*. But, since *SPA* and *SPI* remained unchanged across a range of anesthesia depths, and *SPA* has been shown in MECIII during drug-free sleep, these results should be broadly applicable^16^. On the other hand, a large number of biological factors that we did not consider could modulate our system wide findings. For example, in addition to the direct inputs from the parietal cortex to EC, there is substantial indirect input via the perirhinal and postrhinal cortices that we did not consider^85^. Recent studies show some cortical inhibitory neurons that remain active during the down state, which can alter the nature of cortical UDS^86^. Finally, hippocampus receives EC input and projects back to EC, and EC projects back to the frontal cortices; these connections were not included in our model, but could be studied in the future^87^. Despite this, the simple model was able to predict and match a large amount of experimental observations in a quantitative, cell-by-cell manner. Future studies can build on this approach to study *SPA* and *SPI* during drug-free sleep.

Given the direct and indirect pathways linking the entorhinal region to the hippocampus^40,87^, the decoupling of entorhinal activity from neocortical inputs during *SPA* and *SPI* could contribute to selective removal, strengthening, and weakening of memory traces from the hippocampus during slow-wave sleep, thus improving the signal to noise ratio in the space of memories, thereby improving experimentally observed task-related performance^9^. Our model is sufficiently general and could equally apply to other networks, e.g. parietal-prefrontal network, where persistent activity is seen during working memory tasks^55,88^. Indeed, recent studies of brain activity in humans has shown that functional network connectivity during spontaneous epochs is highly dynamic^89^, and that persistent activity during working memory gates the propagation of activity, and thus information, into the prefrontal network^90^.

In sum, these results demonstrate that during UDS, the rich dynamics of the entire cortico-entorhinal circuit can be captured in a quantitatively precise fashion by a dynamic attractor landscape involving just two biologically important variables: the cortico-entorhinal excitation and the recurrent excitation within the entorhinal cortex. Our model is simple enough to be analytically tractable. Despite the apparent simplicity and with just two parameters, we were able to reproduce nearly a dozen different experimental observations in a quantitatively precise fashion. This provides a strong support for our model to reveal the nature of cortico-entorhinal functional connectivity during slow oscillations *in vivo,* and the differential nature of this connectivity between MECIII vs LECIII. This approach provides a powerful technique to understand the functional connectivity between large networks of neurons *in vivo.*

## Acknowledgements

We thank J. McFarland for doing initial analysis, and noticing SPI, and J. Moore for comments on the manuscript. This work used computational and storage services associated with the Hoffman2 Shared Cluster provided by UCLA Institute for Digital Research and Education’s Research Technology Group.

## Author Contributions

KC did theory, simulations, and data analysis. SB and TGH did experiments. MRM participated and supervised all aspects. KC and MRM wrote the paper, with input from SB.

## Data and Materials Availability

Data and code are available upon reasonable request.

## Supplementary Information

Materials and Methods

Supplementary Fig. 1 to Fig. 11

