## Supplementary Information for "Theory of spontaneous persistent activity and inactivity *in vivo* reveals differential cortico-entorhinal functional connectivity"

### Supplementary Information for Choudhary et. al.

#### Materials and Methods:

##### *A Single E/I Mean Field Model*

We constructed a, mean-field network[45,46] that can support UDS oscillations [20–22,24], which modeled the average activity of two populations of neurons: one excitatory and one inhibitory (Sup. Fig. 1). This average activity level is a dimensionless number whose value ranged between 0 and 1. Since UDS are slow and synchronous oscillations, this level of granularity is sufficient and one does not need to include faster variables like spikes. This greatly reduces the number of free parameters and keeps the equations of motion analytically tractable, with high predictive power. The time evolution of the average excitatory $E(t)$ and inhibitory $I(t)$ activity is governed by:

$$\tau_E \frac{dE}{dt} = -E + \Omega_E(W_{EE}E - W_{EI}I - W_{EA}\,A + \xi + i_E) \quad [1]$$

$$\tau_I \frac{dI}{dt} = -I + \Omega_I(W_{IE}E - W_{II}I + \xi) \quad [2]$$

Where $\tau_{E/I}$ is the time constant of each network ($\tau_E$ = 10 ms, $\tau_I$ =5 ms), consistent with experimental data on the membrane time constants. We first consider the case where there is no external input, so $i_E = 0$. The $\Omega_{E/I}$ response function is a standard threshold-linear function with saturation:

$$\Omega_{E/I}(x) = \begin{cases} 0 & if\ x < \theta \\ g_{E/I}(x - \Theta_{E/I}) & if\ \theta < x < \theta + 1/g_{E/I} \\ 1 & if\ x > \theta + 1/g_{E/I} \end{cases} \quad [3]$$

where $g_{E/I}$ is the slope of the input-output relationship for each neural population ($g_E$ = 6, $g_I$ = 30) and $\theta_{E/I}$ is the threshold input needed for each population ($\theta_E$=0.0517, $\theta_I$=0.2778). For both populations, the inputs are simply the sum of currents from neural populations in the network, given by synaptic weight $W_{XY}$, from population *Y* to population *X*, multiplied by the source activity $E/I$ ($W_{EE}$ = 1, $W_{II}$ = 0.083, $W_{EI}$ = 0.166, $W_{IE}$ = 1.66). There is additional noise current ξ, drawn from a gaussian distribution (mean of 0, std. 0.03) to simulate random fluctuations within the network activity. In all simulations and theoretical analysis, the input remained in the linear regime and never reached saturation. The excitatory population has an additional term to describe its internal, activity-dependent adaptation $A$, with weight $W_{EA}$ = 0.166:

$$\tau_A \frac{dA}{dt} = -A + W_{AE}E \quad [4]$$

Where the time constant $\tau_A = 300\ ms$ is much larger than the time constants of excitation and inhibition, and the modulation due to excitation is $W_{AE}$=1.1. $E = 0, I = 0$ is a steady state of this network, and corresponds to the Down state observed during UDS. Since the adaptation parameter is so slow-varying, we can consider a snapshot of the network at a fixed adaptation $A^*$ and consider the state space of all possible realizations of activity $E/I$. We can solve for the nullclines of each population by setting the derivative of the activity in each population to zero, and solving for $E$ :

$$E = \frac{g_E W_{EI} I + g_E(W_{EA}A^* + \Theta_E)}{g_E W_{EE} - 1} \quad [5]$$

$$E = \frac{(1 + g_I W_{II})I + g_I \Theta_I}{g_I W_{IE}} \quad [6]$$

These are plotted in Fig 1B and Sup. Fig 1-3. In order to have a stable Up state, these two nullclines must intersect at nonzero values for *E* and *I*, which is possible under the condition that

$$\Theta_I > \frac{g_E W_{IE}}{g_E W_{EE} - g_E W_{EA} W_{AE} - 1} \Theta_E \quad [7]$$

$$g_E < \frac{1 + g_I W_{II}}{W_{EE}(1 + g_I W_{II}) - g_I W_{EI} W_{IE}} \quad [8]$$

These conditions are satisfied by our choice of parameters, and ensure that the excitation nullcline is steeper than the inhibition nullcline but has a smaller $E$-intercept, thus ensuring an intersection. We identify this intersection with the neurological Up state, where both excitatory and inhibitory populations exhibit sustained firing. Its coordinates are

$$E = \frac{W_{EI}\Theta_I - \left[W_{II} + \frac{1}{g_I}\right] * \Theta_E}{W_{EI}W_{IE} - \left[W_{EE} - \frac{1}{g_E} - W_{EA}W_{AE}\right] * \left[W_{II} + \frac{1}{g_I}\right]} \quad [9]$$

$$I = \frac{\left[W_{EE} - \frac{1}{g_E} - W_{EA}W_{AE}\right] * \Theta_I - W_{IE}\Theta_E}{W_{EI}W_{IE} - \left[W_{EE} - \frac{1}{g_E} - W_{EA}W_{AE}\right] * \left[W_{II} + \frac{1}{g_I}\right]} \quad [10]$$

A third fixed point is found at $E = 0$ and $I = \frac{g_E(W_{EA}A^* + \Theta_E)}{g_E W_{EE} - 1}$. This point is unstable and lies on the separatrix, which marks the boundary between two regions of stability. Thus, one can imagine that the network sits in a potential landscape with two minima and one energy barrier in the middle [91].

The local stability of the Up state can be found by linearizing the differential equations 1-2 about the Up state fixed point, and ensuring that the eigenvalues of the matrix of coefficients have a negative real part, signifying that fluctuations will exponentially decrease [21,48]. For our 2D matrix, this is equivalent to imposing that the determinant of the coefficients matrix is positive and the trace is negative. These conditions yield the following relations between connectivity, time-scale, and gain:

$$\left[W_{II} + \frac{1}{g_I}\right] * \left[W_{EE} + \frac{1}{g_E}\right] < W_{EI} * W_{IE} \quad [11]$$

$$\tau_I * (g_E W_{EE} + 1) < \tau_E * (g_I W_{II} + 1) \quad [12]$$

These conditions are satisfied by our choice of parameters.

The global stability of each fixed point is inversely related to the distance of the fixed point from the unstable separatrix. The closer each stable fixed point (the Up or Down state) is to the separatrix, the less relatively stable that fixed point becomes, since random noise has a higher chance of kicking the network over the boundary. Notably, the variable $A^*$ is simply an additive constant to the excitation nullcline, the dynamics of which determines the positions of intersection for both the stable Up state as well as the separatrix. As the network remains in the Up state, the adaptation variable increases, effectively shifting the excitation nullcline up. This not only decreases the overall firing rate in the Up state by shifting the fixed point, it also stabilizes the Up state by bringing the separatrix closer to the Up state. A kick from random noise eventually forces the network to transition into the down state. Here, adaptation recovers back to zero, shifting the excitation nullcline down, thereby bringing the separatrix closer to the Down state fixed point; eventually, a noisy kick forces the network into the Up state, where the cycle repeats.

***Coupled Networks: Persistent Activity/Inactivity***

To model spontaneous persistent activity, we consider two identical networks, each described by the above equations. Further, the afferent network provides a weak excitatory input $W_{EXT}$ into the excitatory population of the efferent network. We use $W_{EXT}$ to refer to this synaptic weight from the afferent to the efferent network and $W_{INT}$ to refer to the internal excitatory-excitatory weight in the efferent network ($W_{EE}$ in equations 1-12). Effectively, this means that for the efferent network there is an external input into the excitatory population $i_E(t) = W_{EXT} \cdot E_A(t)$, where $E_A(t)$ is the activity of the excitatory population in the afferent network. If the connection strength $W_{EXT}$ between the afferent and efferent excitatory populations is sufficiently strong, the two networks UDS oscillations phase lock, as the transitions between states in the efferent network are no longer due to independent noise but the timed increase and decrease in input coming from the afferent network. Similar to previous results, the connection strength $W_{EXT}$ must be about an order of magnitude smaller than the internal connections $W_{INT}$ in order to show desynchronization [21].

What happens when both networks are in the Up state and the afferent input transitions into the Down state? This cuts off afferent input $W_{EXT}$, immediately shifting the efferent excitation nullcline to the left, thereby destabilizing the efferent Up state. The efferent network can either remain in the Up state through its own recurrent excitation or follow the afferent and transition into the Down state. Instances when the efferent remains in the Up state are termed "spontaneous persistent activity (*SPA*)." It follows from the stability arguments outlined earlier that by increasing the distance between the Up state fixed point and the separatrix, one can increase the stability of the Up state, thereby increasing the probability that the network will display *SPA*. If we refer to the excitation nullcline equation [Eq. 5], we see that increasing $W_{INT}$ ($W_{EE}$ in Eq. 1-12) results in the downward scaling of the nullcline: the Up state fixed point shifts to the right, and the unstable fixed point shifts downward. Combined, this has the effect of increasing the stability of the Up state, thus leading to higher probability of *SPA*.

The converse scenario applies for the Down state, where both networks are in the Down state and the afferent transitions into the Up state. This suddenly increases the input, shifting the excitation nullcline down, destabilizing the Down state. The efferent network can either follow into the Up state or remain in the Down state; the latter case we term "spontaneous persistent inactivity" (*SPI*). The size of downward shift due to the incoming current from the $W_{EXT}$ synapse has a direct consequence on the stability of the Down state: the larger the weight, the more the shift, and thus the more destabilized the DOWN state becomes. Thus, decreasing the synaptic weight $W_{EXT}$ increases the probability of *SPI*. Indeed, simulations where we modulated both $W_{INT}$ and $W_{EXT}$ confirmed our hypothesis on the dependence of persistent activity and inactivity on these two variables (Fig 1B). All simulations were performed using MATLAB on the UCLA Hoffman2 Computing Cluster with time-step 0.2 ms using the Runge-Kutta method.

***Animals, Surgery, and Histology***

Methods were similar to those described previously [16]. Briefly, data were obtained from 136 C57BL6 mice aged postnatal day (p)25-43 (p32±1) weighing 12-21 g (17.5±0.4 g). Mice were anesthetized with urethane (1.64±0.03 g urethane / kg body weight intraperitoneal). Body temperature was maintained at 37℃ with the help of a heating blanket. The animals were head-fixed in a stereotaxic apparatus and the skulls exposed. A metal plate was attached to the skull and a chamber formed with dental acrylic, which was filled with warm cerebrospinal fluid. Two 1-mm diameter holes, one for the LFP recordings and one for the whole-cell recordings, were drilled over the left hemisphere and the underlying dura mater was removed.

After electrophysiological recordings, mice were euthanized by transcardial perfusion with 0.1 M phosphate buffer, followed by 4% Paraformaldehyde solution, and 150-200 µm thick brain sections were processed with the avidin-biotin-peroxidase method. Sometimes, a subsequent Nissl stain was applied before embedding. Visualization of biocytin filled neurons allowed for the determination of cell type and recording site. Unidentified neurons were excluded from analysis. All experimental procedures were carried out according to the animal welfare guidelines of the Max-Plank-Society.

##### *Electrophysiology and Data Acquisition*

Local field potentials (LFPs) were recorded with an 8 site single-shank multisite probe (NeuroNexus Technologies). LFP from layer 2/3 of posterior parietal cortex (2 mm posterior to bregma, 1.5 mm lateral) was used to characterize neocortical Up-Down states. *In vivo* intracellular membrane potential ($V_m$) was recorded in whole-cell configuration by using borosilicate glass patch pipettes with DC resistances of 4-8 MΩ and filled with a solution containing 135 mM Potassium Gluconate, 10 mM HEPES, 4 mM Potassium Chloride, 10 mM Phosphocreatine, 4 mM MgATP, 0.3 mM $Na_3GTP$ (adjusted to pH 7.2 with KOH), and 0.2% biocytin for subsequent histological identification. Whole-cell recording configuration was achieved as described previously[92]. Relative to bregma, the anteroposterior (AP), mediolateral (ML) and dorsoventral (DV) coordinates of the craniotomies for the $V_m$ recordings were made around -4.5 mm AP and 4 mm ML for MEC; -3.5 to -4 mm AP, 4.5 mm ML and 4 mm DV for LEC; -1.5 to -2 mm AP and 1 mm ML for parietal cortex; 1 to 1.5 mm AP and 1 mm ML for frontal cortex; 2 to 3 mm AP and 0.5 to 1 mm ML for prefrontal cortex.

The average initial series resistance was 46 MΩ, and $V_m$ values were corrected for the estimated junction potential of approximately +7 mV.

The $V_m$ was acquired by Axoclamp-2B (Axon Instruments) and fed into a Lynx-8 amplifier (Neuralynx). The $V_m$ and LFP were recorded by an HS16 preamplifier (Neuralynx) for about 20-40 minutes. The complete recording was used for subsequent statistical analysis. The LFP were sampled at 2 kHz, low-pass filtered below 475 Hz, and amplified 1000-5000 times. The membrane potential was low-pass filtered below 9 kHz, sampled at 32 kHz, and amplified 50-150 times. Simultaneously, the DC value of $V_m$ was recorded by an ITC18 interface (Instrutech) under the control of Pulse software (Heka) or by a Micro1401 with Spike2 software (CED). Some of these DC-coupled data were recorded in discontinuous sweeps of 7 or 10 s, separated by 5 or 2 s, respectively. Data from previous work [16] was supplemented with additional recordings from MECIII and LECIII. The parietal, frontal, and prefrontal $V_m$ measurements are entirely new.

##### *Data preprocessing*

All analysis was restricted to subthreshold fluctuations in the membrane potential by removing spikes as follows. The temporal derivative of the bandpass-filtered (100 Hz - 8 kHz) membrane potential signal was computed, and times when this derivative exceeded 10 standard deviations above the mean were taken as spike times. Spike waveforms were then removed by replacing 3 ms of data following the onset of each spike by linear interpolation of adjacent values. To remove the 50 Hz mains hum and its many harmonics, 8-pole bandstop filters were used at 45-55 Hz, 95-105 Hz, 145-155 Hz, 195-205 Hz, 245-255 Hz, and 295-305 Hz.

***Explicit-duration Hidden Markov Model detection of Up and Down states.***

Synchronized epochs, wherein the LFP and cortical $V_m$ undergoes synchronous transitions of Up and Down states (UDS), were selected by locating and eliminating periods of data with desynchronized activity where UDS are absent. Previously outlined methods[50] were closely followed. Briefly, the spectrogram of the signal was computed in 15-s overlapping windows using multi-taper methods (Chronux Matlab toolbox) with a time-bandwidth product of 4, and seven tapers. The maximum log power in the range of 0.05-2 Hz and the integral of the log power in the 4-40 Hz range we then used to locate and remove desynchronized epochs in the data.

The remaining data exhibited UDS. UDS of both membrane potential and neocortical LFP were classified using two state explicit-duration hidden Markov models (EDHMMs). The $V_m$ and LFP were first filtered in the low frequency (0.05-2 Hz) range, and a gaussian observation EDHMM was fit to the filtered signal, with inverse gaussian models of the state duration distributions. The means of the state-conditional gaussians were slowly varying functions of time, where the parameters were estimated over a 50 second window length. We found the maximum likelihood parameter estimates of the EDHMM, and computed the resulting "Viterbi" sequence, which was used to define UDS oscillations.

***Assignment of corresponding neocortical-entorhinal state transitions.***

Given two UDS sequences, one for the neocortical LFP (the afferent network in the simulation) and one for the entorhinal $V_m$ (efferent network), the fine temporal relationships and quantized duration was calculated by first assigning each Up/Down state in the $V_m$ to its corresponding set of trigger states in the LFP. This was done through a greedy search algorithm, where in each iteration of the algorithm, the Up/Down state initiations were linked to the closest Up/Down state initiations in the corresponding LFP. Note that this does not guarantee a one-to-one mapping from $V_m$ states to LFP states; those $V_m$ states which map onto more than one LFP state are termed "persistent." The quantized duration of a $V_m$ state was calculated as the number of total LFP states (both Up and Down) that would fit inside a particular $V_m$ state, with each Up and Down state in the cycle contributing to 0.5 units of time (Sup. Fig 10).

***Model Fitting***

The $W_{EXT}$ - $W_{INT}$ parameter space was divided into a 100-100 grid, and each point was taken as the input into 5 independent simulations of length 1000s. For each experiment, we calculated the "distance" between the experimental data *SPA*/*SPI* level and each simulated *SPA*/*SPI* level. Let $\phi_{SPA/SPI}$ denote the proportion of efferent states which were classified as *SPA*/*SPI* (see above) in the experiment, and $\xi_{SPA/SPI}$ denote the proportion in a given simulation. The distance between the experiment and any particular simulation is then given by $d = \sqrt{(\phi_{SPA} - \xi_{SPA})^2 + (\phi_{SPI} - \xi_{SPI})^2}$. The simulation with the minimum distance to the experiment was chosen as the best fit (Fig 3A). One could also take $\phi$ and $\xi$ to denote the proportion of afferent LFP states which were classified as 'skipped' by the efferent network, and use this for the distance metric. Results of this fit are shown in Sup. Fig 7, and are virtually identical to the procedure used in Fig 3A of the main text.

***Statistics and hypothesis testing***

Central Tendencies and variability is reported as mean plus/minus standard deviation, unless otherwise noted. All hypothesis tests were performed using two-sided nonparametric Wilcoxon rank-sum tests for equal medians. Wilcoxon signed rank tests were used for paired comparisons or one-sample tests. Correlations were computed using Spearman's rank correlation coefficient. A p-value of less than 0.05 was used for statistical significance.

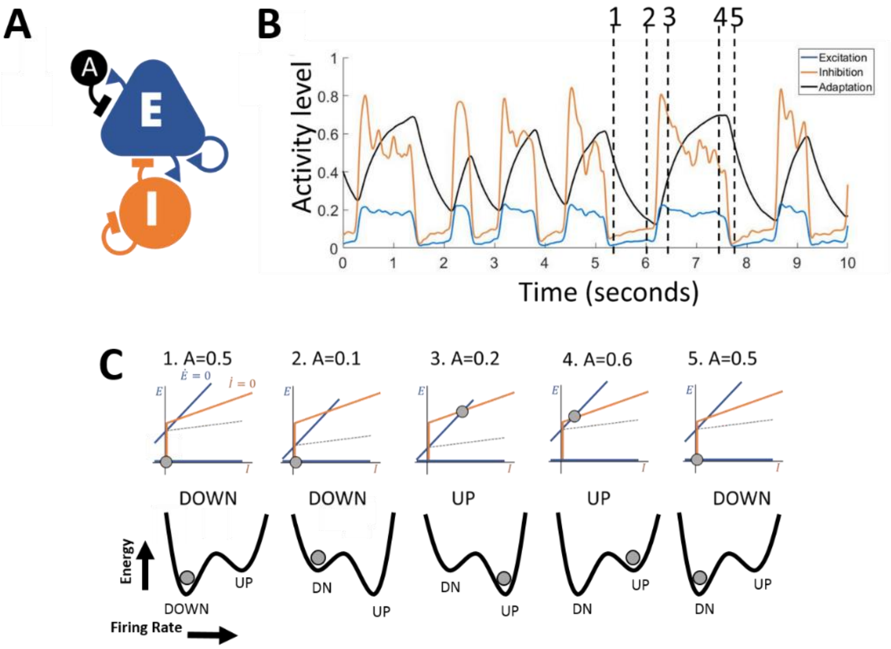


**Supplementary Fig. 1: A single network of inhibitory and adaptation mediated excitatory populations can produce Up-Down state oscillations**. **A)** Excitatory and inhibitory populations are recurrently connected, and their activities are quantified by variables *E* and *I*, which vary between 0 and 1 and represent the proportion of active (spiking) neurons in the population. Only the excitatory population has activity dependent adaptation α, which evolves with time constant $\tau_\alpha$=500s. **B)** A sample trace of *E, I,* and α. **C)** Top row: The time-evolution of the network for one complete UDS cycle (time points in B) can be visualized on the *E-I* phase space. Nullclines for the *E* and *I* variables are plotted in blue and orange, respectively, and denote where the time derivative for that particular variable ($\dot{E}, \dot{I}$) is zero. Intersections of the nullclines denote equilibrium points where both $\dot{E} = 0$ and $\dot{I} = 0$. There are two stable points, corresponding to the Up state (*E,I*>0) and the Down state (*E,I*=0). These two attractor points form basins of attraction in the *E-I* plane, with a separatrix (dashed line) denoting the boundary. Under noise, the relative stability of each point is inversely proportional to its distance from this separatrix, and the entire system can be viewed as an energy landscape, with each stable point as an energy minimum (bottom row). An increase in α corresponds to an upwards translation of the $\dot{E}$ nullcline, making the Up state less stable; a decrease in α corresponds to a downward shift, making the Down state less stable.

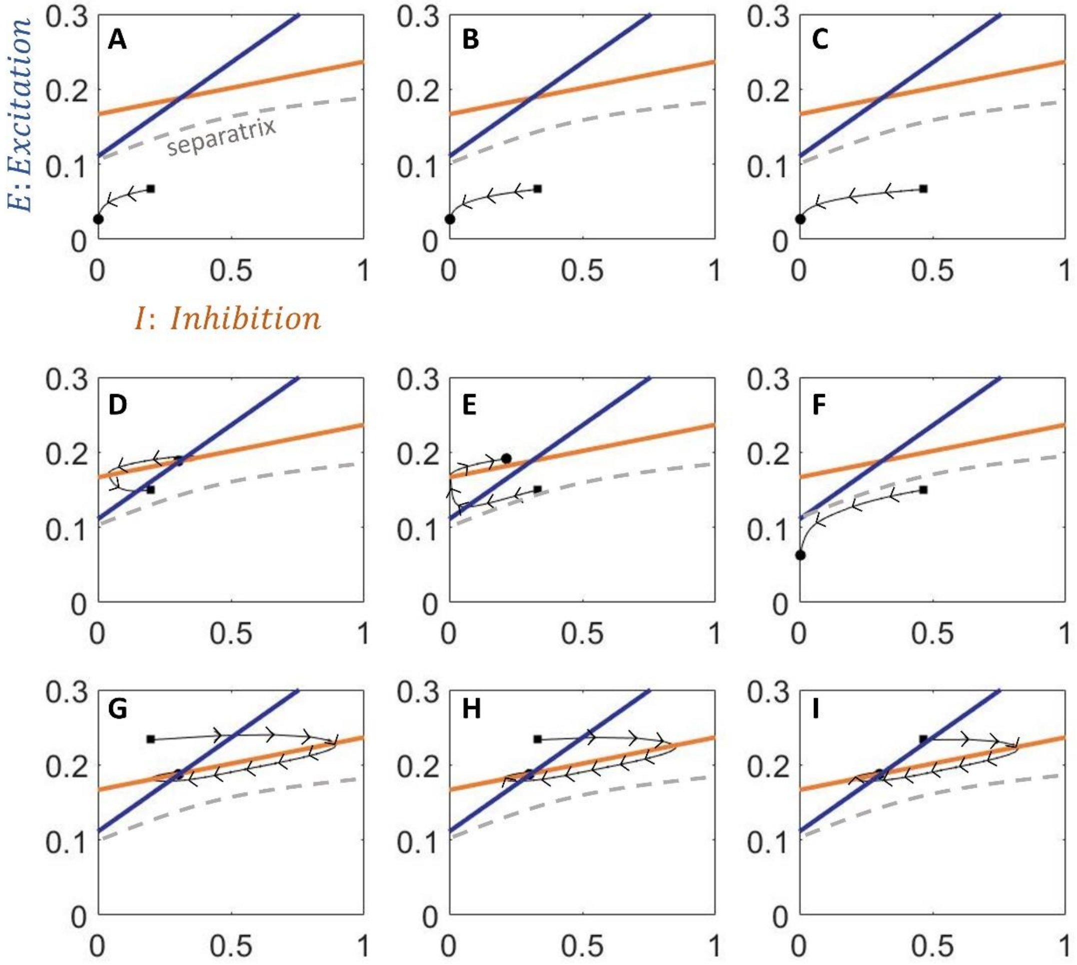


**Supplementary Fig. 2: The mean field equations define an attractor landscape in the *E-I* coordinate plane.** The mean field model defines two discrete attractors in the *E-I* coordinate plane. Here we trace the evolution of a single network in the $(E, I)$ coordinate plane from various initial conditions (denoted by the black square) to the final condition (denoted by the circle) in the absence of noise and for fixed adaptation level (α=0.5). The excitation and inhibition null clines are depicted (solid colored lines) along with the separatrix (dotted line). Conditions A, B, C, and F end in the 'Down' state fixed point, while the others end in the 'Up' state fixed point.

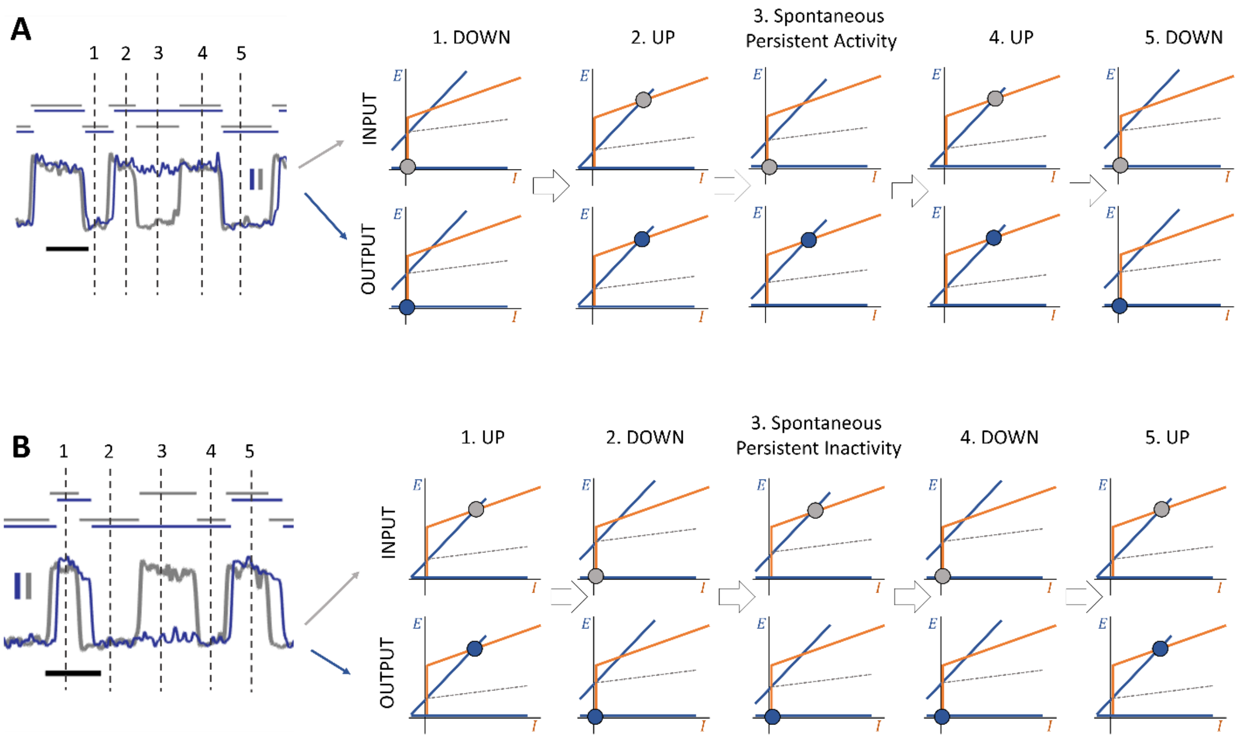


**Supplementary Fig. 3: *SPA* and *SPI* occur when the efferent network persists in its current state and does not follow a state transition in the afferent network. A)** Left: *SPA* occurs when the efferent network (blue) persists in the Up state while the afferent (gray) undergoes a complete Down state. The black scale bar represents 1 sec in time, and the blue and gray scale bars correspond to activity of 0.1. Right: Each network lives in its own *E*-*I* coordinate plane. A sudden decrease (from 2->3) in the afferent input (an Up-Down transition) translates the efferent output *E* nullcline upward, destabilizing the Up state and inducing a synchronous transition to the Down state. If the destabilization is not enough (because the decrease was not large, i.e. the afferent Down state has high activity), the efferent network can persist in the Up state on its own, resulting in *SPA*. **B)** Similar to A, but showing *SPI*, which occurs when the efferent network persists in the Down state while the afferent undergoes a complete Up state. A sudden increase in the afferent input (a Down-Up transition) translates the efferent *E* nullcline downwards, destabilizing the Down state an inducing a transition to the Up state. Again, if the destabilization is not enough (because the afferent Up state has low activity), the efferent network can persist in the Down state, resulting in *SPI*. Scale bars show time (horizontal black, 1 second) and the amplitude in afferent (vertical gray, 0.1) and efferent (vertical blue, 0.1) network activity.

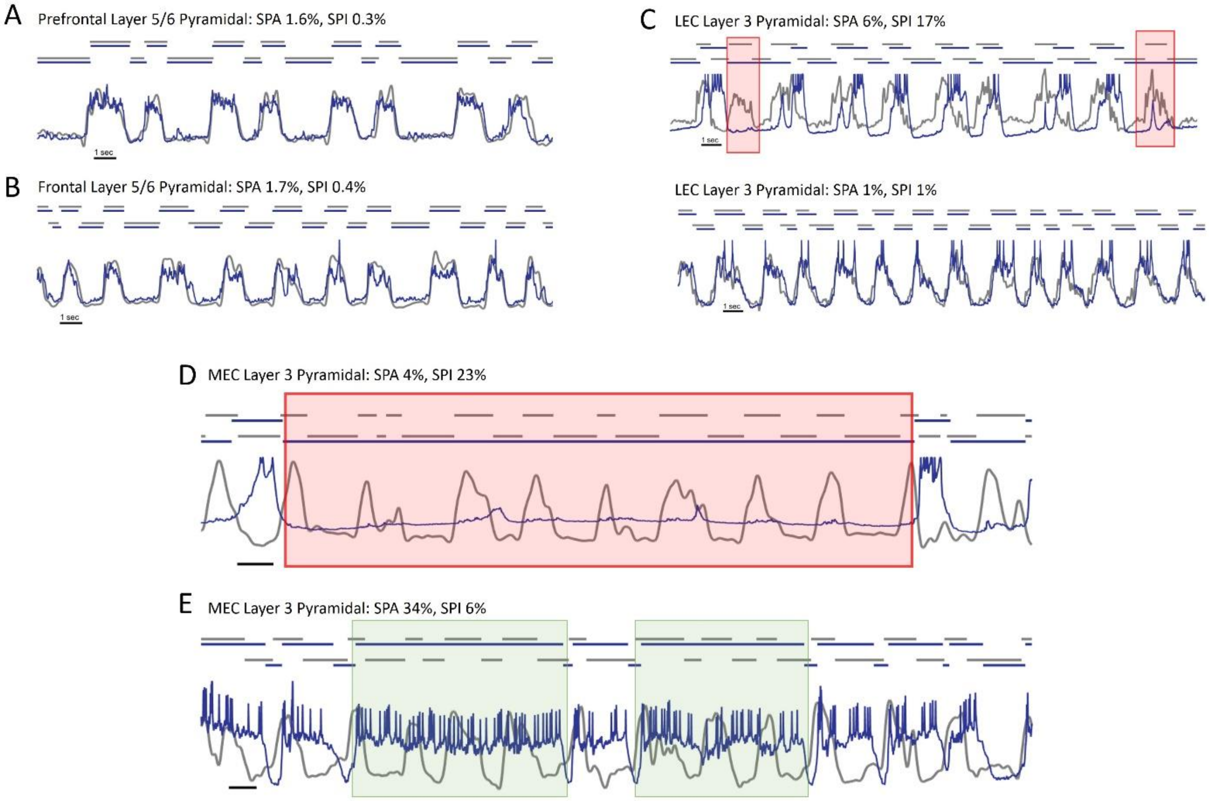


**Supplementary Fig. 4: Additional examples of experimental traces A)** Example trace of the $V_m$ (blue) of a prefrontal cortex layer 5/6 pyramidal neuron, along with simultaneously recorded neocortical LFP (gray). Black scale bar at the bottom shows one second time interval, and the detected Up-Down state sequence is shown above for each trace. **B)** Example trace from frontal cortex layer 5/6 pyramidal neuron. Both neurons (in A and B) exhibited phase-locked UDS to the neocortical LFP. **C)** Two examples of LECIII pyramidal neurons, one showing heightened levels of persistent inactivity (top, red boxes), and the other showing complete phase locking (bottom). **D)** An example of MECIII neuron showing extremely long persistent inactivity state (red box), lasting over 17.2 seconds. **E)** An example of MECIII cell showing extremely long persistent activity states (green boxes), lasting 8 and 6 seconds.

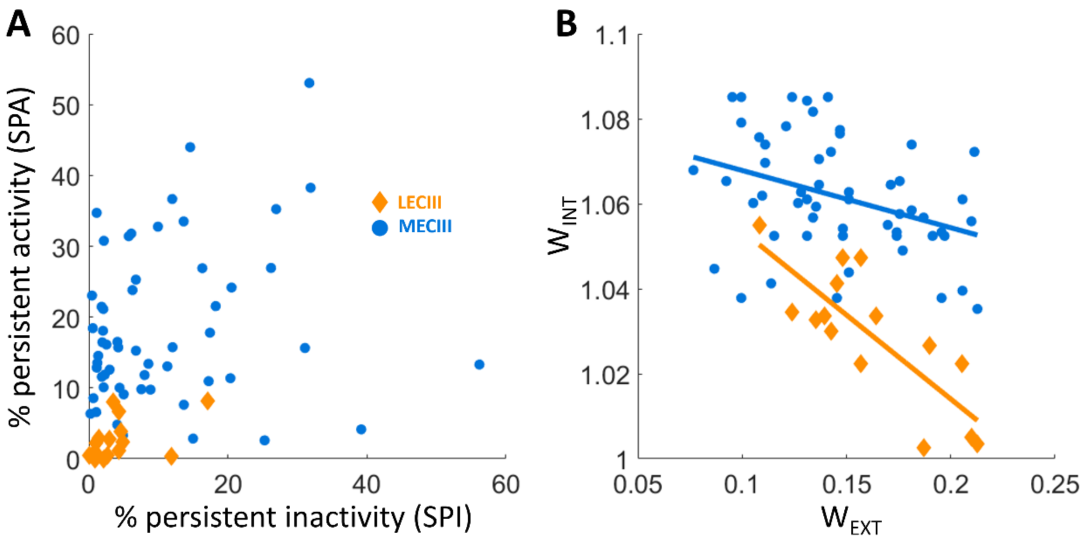


**Supplementary Fig. 5: The amount of *SPA* in an entorhinal cell was not significantly correlated with the amount of *SPI*, but fitted parameters $W_{INT}$ and $W_{EXT}$ are significantly anti-correlated. A)** Both MECIII and LECIII cells showed varied levels of *SPA* and *SPI* in experiment, but there was no significant correlation between the average *SPA* and *SPI* exhibited by a cell (MECIII: r=0.18, $p>10^{-1}$; LECIII: r=0.49, $p>10^{-1}$). **B)** On the other hand, fitted parameters $W_{INT}$ and $W_{EXT}$ were significantly anti-correlated within both MECIII (r=-0.36, $p<10^{-2}$) and LECIII (r=-0.80, $p<10^{-3}$).

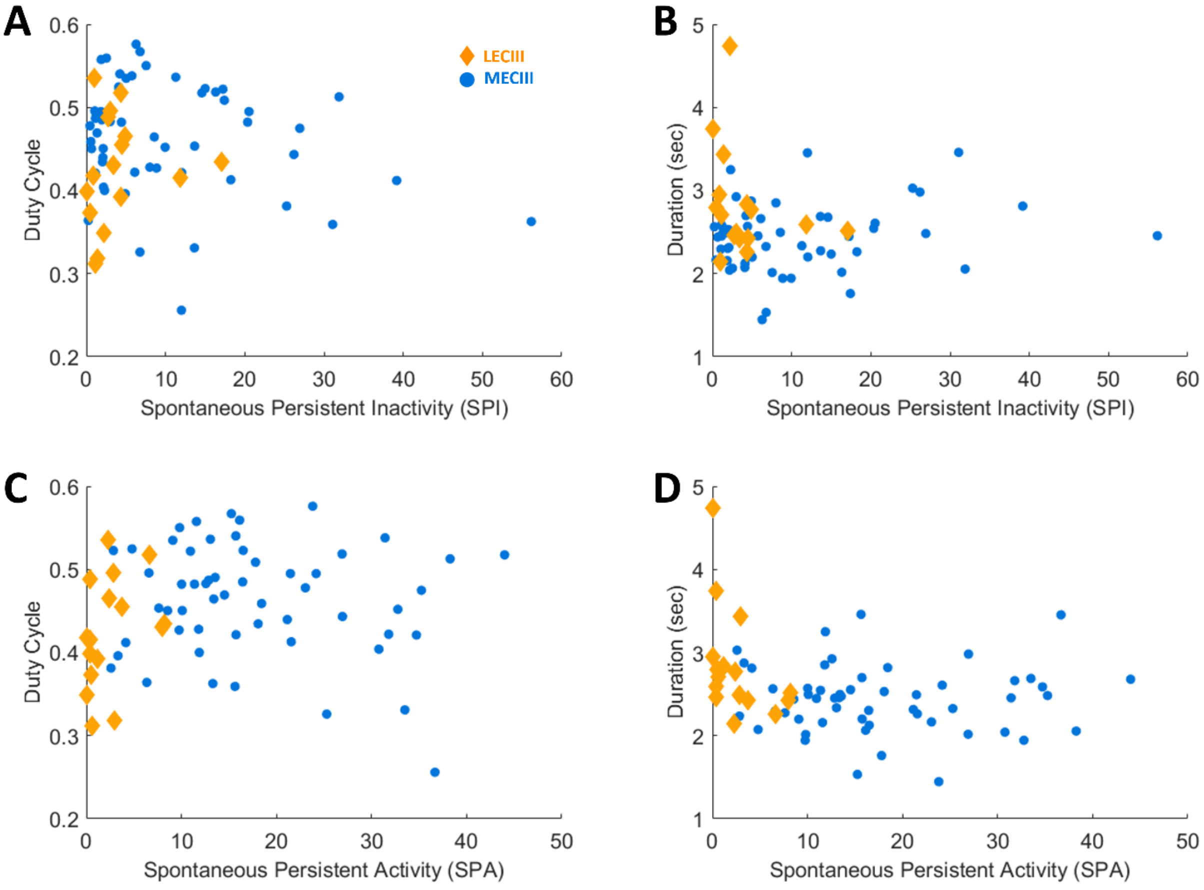


**Supplementary Fig. 6: The prevalence of *SPA* or *SPI* in an experiment was independent of the depth of anesthesia.** The depth of anesthesia varies from one experiment to another, and deeper anesthesia produces UDS with longer duration cycles and smaller duty cycle (proportion of the cycle spent in the Up state). Neocortical UDS statistics were used to quantify depth of anesthesia of the animal. **A)** *SPI* prevalence was not significantly correlated with neocortical duty cycle for both LECIII neurons (yellow: r=0.15, p>0.1) and MECIII neurons (blue: r=-0.24, p>0.05). **B)** *SPI* was not significantly correlated with mean duration of neocortical UDS cycles in the experiment (LECIII: r=-0.27, p>0.3; MECIII: r=0.17, p>0.2). **C)** Conversely, *SPA* was not significantly correlated with neocortical UDS duty cycle (LECIII: r=0.38, p>0.3; MECIII: r=-0.12, p>0.3) and **D)** not correlated with mean duration of UDS (LECIII: r=-0.43, p>0.05; MECIII: r=-0.01, p>0.9).

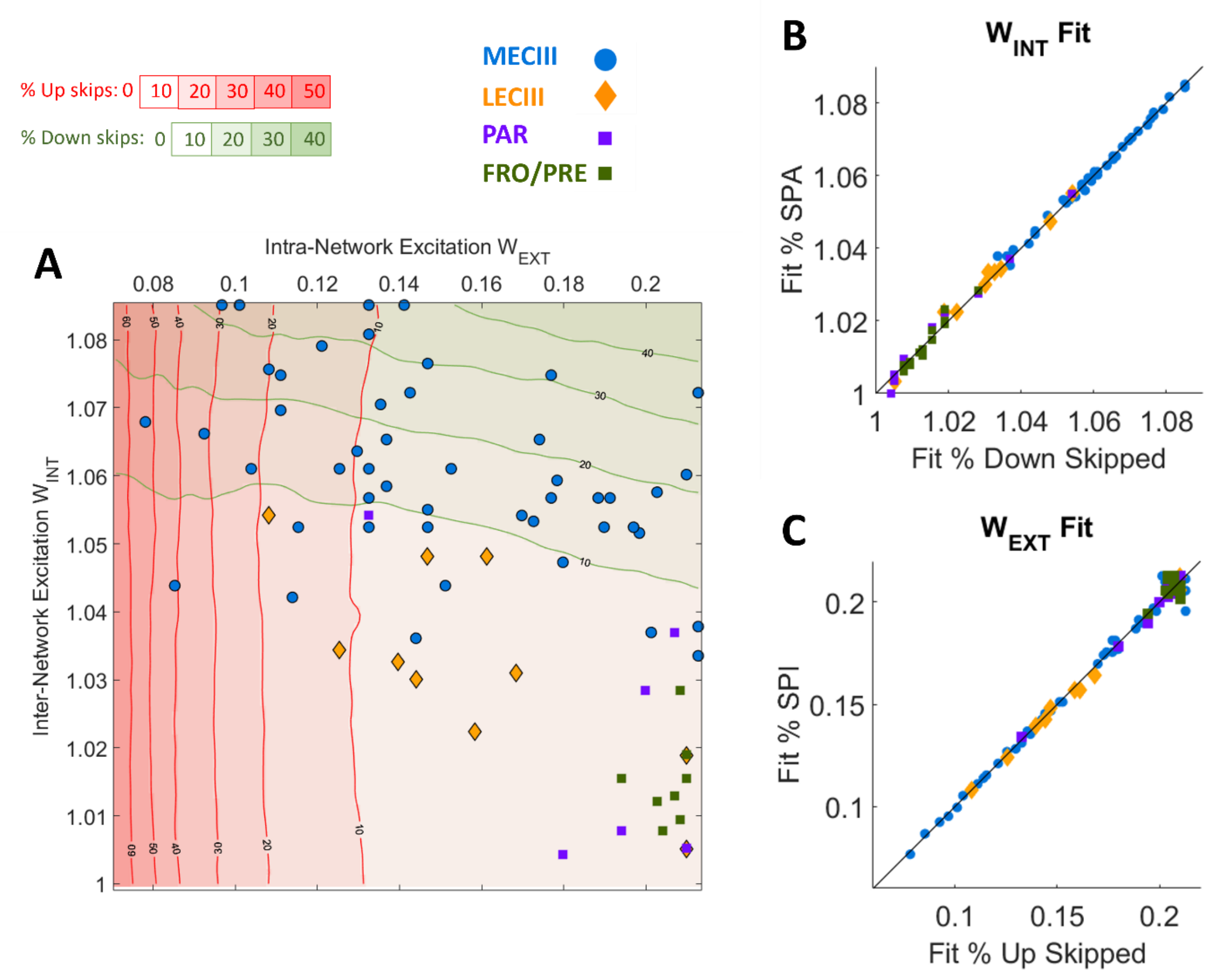


**Supplementary Fig. 7: Experimental fits to the model were robust over different fitting procedures.** In the main text (see Fig 2A), the time-averaged prevalence of *SPA* and *SPI* in the efferent states was used to match simulations to the experiment. **A)** Another method of fitting experimental data to the model is to use the time averaged proportion of *afferent* states that are "skipped" by the efferent network (skipping afferent Up states leads to *SPI*, while skipping afferent Down states leads to *SPA*). Each cell is matched to the simulation space using this metric. The symbols used for each brain region are the same as in Fig 2A. **B)** Comparing the fits to $W_{INT}$ resulting from this matching scheme (x-axis) to the %*SPA*/*SPI* scheme used in the main text (y-axis) reveals that both methods give extremely close values (diagonal line is perfect agreement). **C)** Comparing the fits to $W_{EXT}$ also shows stark agreement.

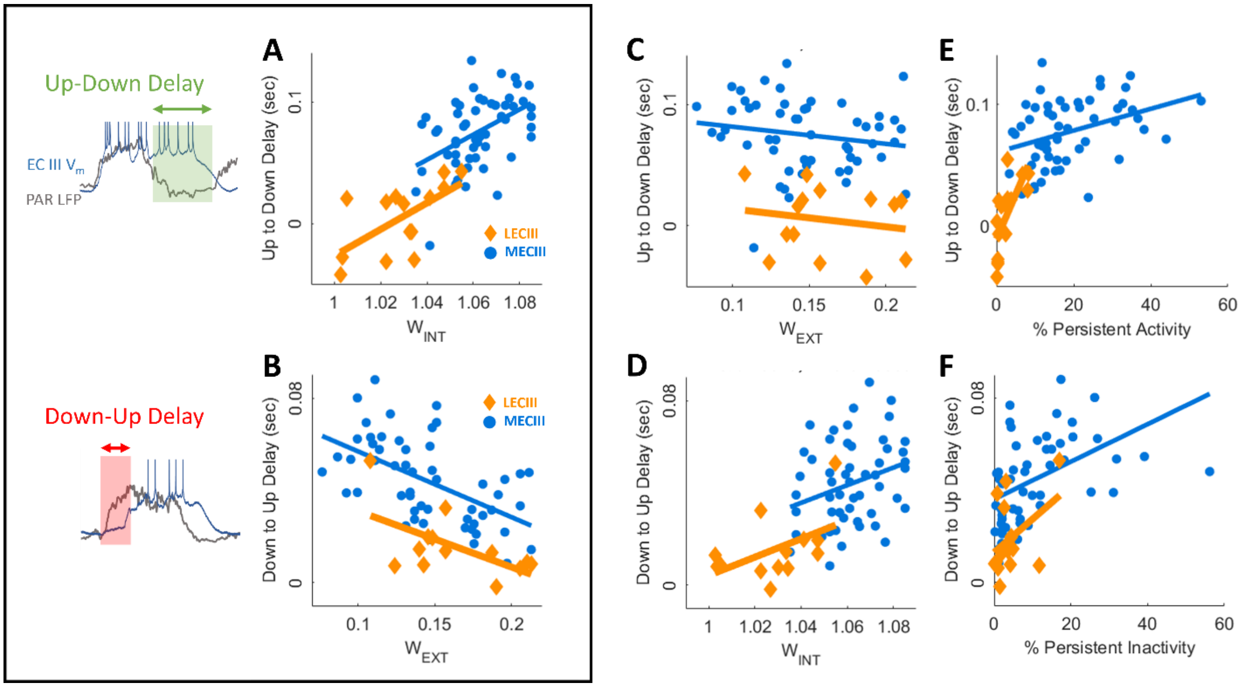


**Supplementary Fig. 8: Up to Down transition lag between afferent neocortical LFP and ECIII $V_m$ is most correlated with recurrent excitation $W_{INT}$, while the Down to Up transition lag is most correlated with external input strength $W_{EXT}$. A)** Same as main text Fig 2E. The internal recurrent excitation $W_{INT}$ increases the stability of efferent network Up state, leading to higher Up-Down delays w.r.t. neocortical LFP (MECIII: blue, r=0.47, $p<10^{-3}$; LECIII: yellow, r=0.63, $p<10^{-2}$). All delays are reported in units of mean UDS duration. **B)** Same as Fig 2G. $W_{EXT}$ is significantly correlated with the Down to Up Delay (MECIII: r=-0.56, $p<10^{-5}$; LECIII: yellow, r=-0.60, $p<10^{-2}$). **C)** $W_{EXT}$ is negatively correlated with the Up-Down delay, but this is not significant (MECIII: r=-0.17, $p>10^{-1}$; LECIII: yellow, r=-0.17, $p>10^{-1}$). **D)** $W_{INT}$ is positively correlated with the Down-Up delay, but this is significant for the MECIII population (blue, r=0.29, $p<0.05$) and not significant for the LECIII population (yellow, r=0.49, $p>10^{-1}$). **E)** Previous studies [16] reported a strong correlation between the amount of *SPA* in a neuron and the Up-Down delay. This is reproduced here. Both MECIII and LECIII populations show significant positive correlation (MECIII: r=0.36, $p<10^{-2}$; LECIII: yellow, r=0.70, $p<10^{-2}$). **F)** Conversely, the amount of *SPI* in a neuron was correlated with the Down-Up delay. The MECIII population showed significant positive correlation (r=0.48, $p<10^{-3}$), while LECIII showed positive but not significant correlation (r=0.42, $p>10^{-1}$).

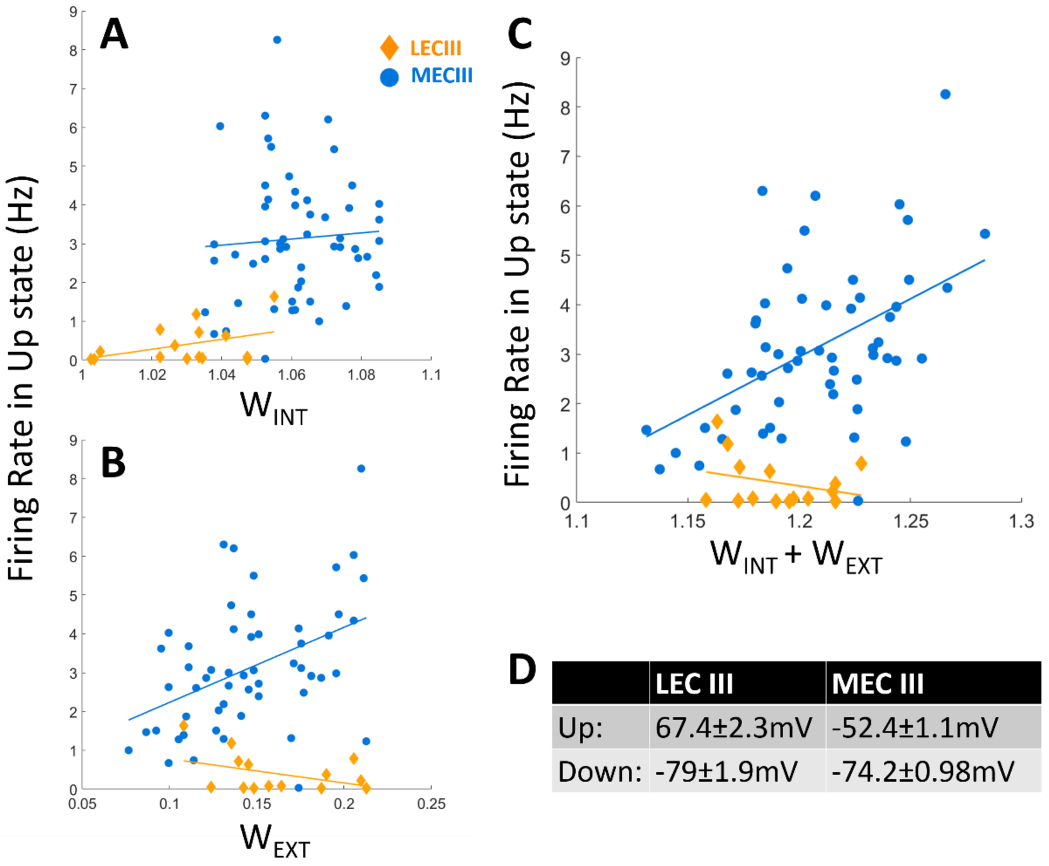


| | LEC III | MEC III |
|---|---|---|
| Up: | 67.4±2.3mV | -52.4±1.1mV |
| Down: | -79±1.9mV | -74.2±0.98mV |

**Supplementary Fig. 9: Firing rate of MECIII but not LECIII cells was significantly correlated to inferred connectivity parameters. A)** The average firing rate of a neuron during the Up state was not significantly correlated with our estimate for the recurrent excitation $W_{INT}$ for either the MECIII (blue: r=0.07, p>0.5) or LECIII (yellow: r=0.42, p>$10^{-1}$) populations. **B)** The average firing rate in the Up state was significantly correlated with our estimate for the external input $W_{EXT}$ for only the MECIII population (r=0.43, p<$10^{-3}$) and not for the LECIII population (r=-0.4, p>$10^{-1}$). **C)** To find the total excitatory input into a neuron, we used the sum of our estimates for the recurrent and external excitations $W_{EXT} + W_{INT}$. The MECIII population showed significant positive correlation (r=0.49, p<$10^{-3}$) while the LECIII population did not show significant correlation (r=-0.29, p>$10^{-1}$). **D)** MECIII neurons were significantly more depolarized than LECIII neurons, during both Up and Down states (p<$10^{-7}$).

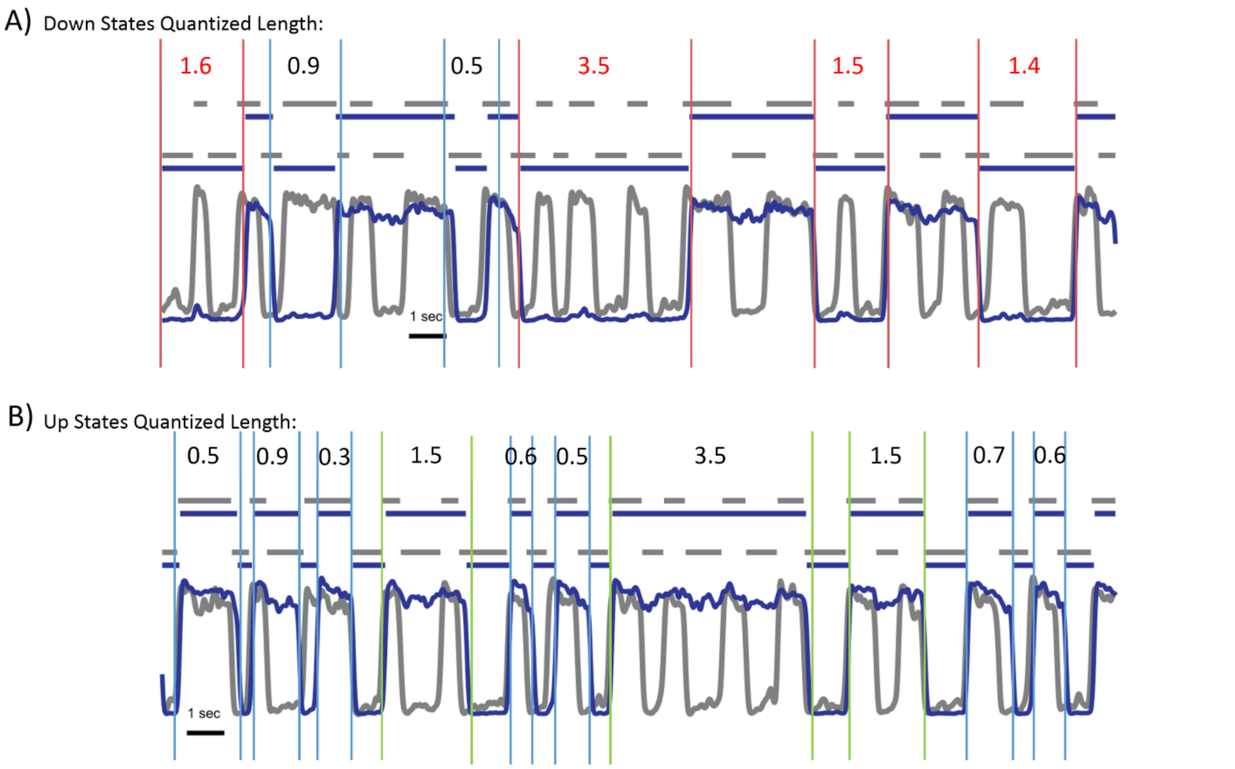


**Supplementary Fig. 10: Counting membrane potential state durations in terms of neocortical UDS cycles. A)** 'Down' states in the membrane potential (blue trace) are first aligned with the nearest neocortical 'Down' state, and the number of LFP 'Down' and 'Up' states this particular membrane potential 'Down' state lasts is counted. Each 'Down' and 'Up' state receives a time length of 0.5 units, a full UDS cycle is thus 1-unit long. The 'Down' states in red represent those that are considered "spontaneous persistent inactivity," since they last for longer than one full UDS cycle. **B)** Same as A, but for 'Up' states. Those 'Up' states that were identified as persistent activity are highlighted in green.

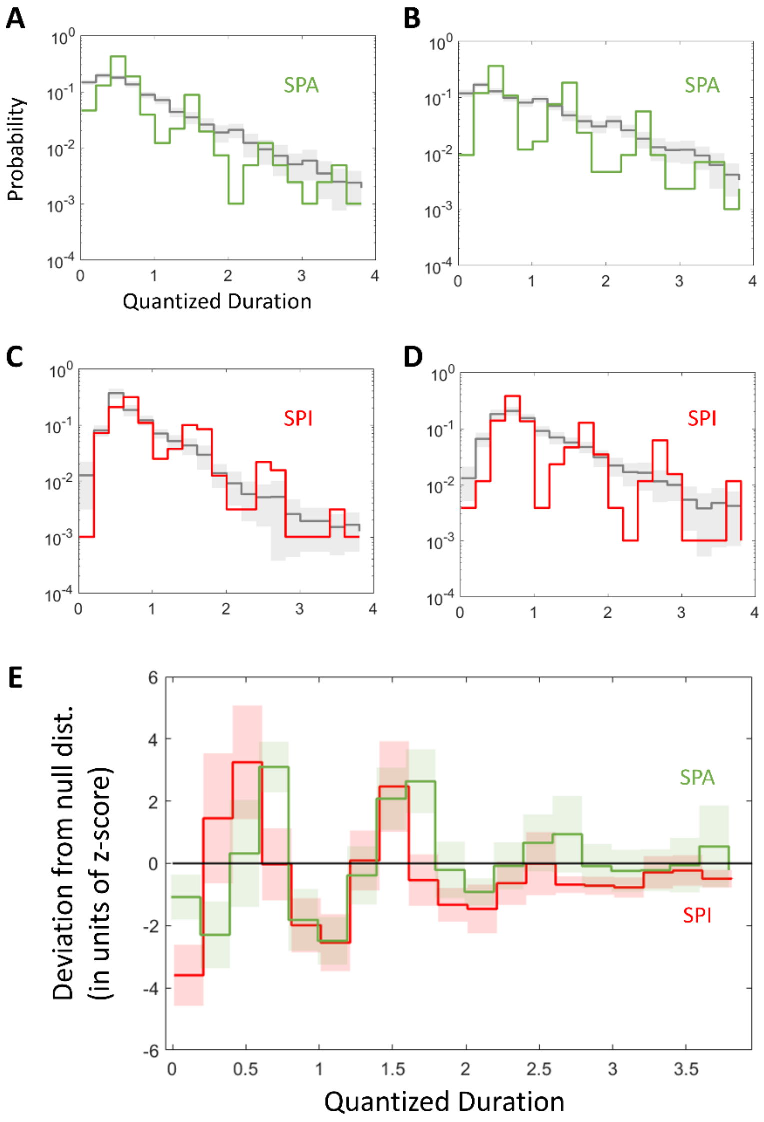


**Supplementary Fig. 11: Experimental distributions of efferent state length in units of afferent UDS was significantly quantized compared to bootstrap shifted null distributions. A)** The distribution of efferent Up state lengths in terms of afferent UDS in the experiment (green) was compared with the same distribution resulting from a random shift of the efferent data (gray). The standard deviation (gray shaded area) of the null distribution was obtained by calculating the distribution from 60 random shifts. The peaks and troughs of the real distribution were significantly different from the null. **B)** Another example of efferent *SPA* quantization, with corresponding null distribution. **C)** An example of efferent Down state lengths (red) in terms of afferent UDS also shows significant quantization, different from the null distribution (gray). **D)** Another example of significant quantization of *SPI*. **E)** Taking the difference between the real and null distributions from each experiment and normalizing by the standard deviation in each bin shows that the first trough (at quantized duration (QD) =1) and the second peak (at QD=1.5) for both *SPA* and *SPI* is significantly different (*SPA*: first trough $p<10^{-3}$, second peak $p<10^{-3}$; *SPI*: first trough $p<10^{-3}$, second peak $p<10^{-2}$) from the null distribution at the population level. The SPA QD peaks were shifted to the right of the SPI peaks, but the troughs occurred at the same QD. This reflects the fact that for MECIII neurons the Up-Down latency is significantly larger than the Down-Up latency, and that Up state termination is internally determined, while Up state initiation is affected strongly by external input.
